# Circadian rhythm disruptions associated with opioid use disorder in the synaptic proteomes of the human dorsolateral prefrontal cortex and nucleus accumbens

**DOI:** 10.1101/2023.04.07.536056

**Authors:** Stephanie Puig, Xiangning Xue, Ryan Salisbury, Micah A. Shelton, Sam-Moon Kim, Mariah A. Hildebrand, Jill R. Glausier, Zachary Freyberg, George C. Tseng, Anastasia K. Yocum, David A. Lewis, Marianne L. Seney, Matthew L. MacDonald, Ryan W. Logan

**Author notes:** **Co-corresponding authors** Ryan W. Logan, PhD 364 Plantation Street, LRB Office 611 Worcester, MA 01605; Matthew L. MacDonald 203 Lothrop St, Thomas E. Starzl Biomedical Science Tower, Room W1655 Pittsburgh, PA 15213.

## Abstract

Opioid craving and relapse vulnerability is associated with severe and persistent sleep and circadian rhythm disruptions. Understanding the neurobiological underpinnings of circadian rhythms and opioid use disorder (OUD) may prove valuable for developing new treatments for opioid addiction. Previous work indicated molecular rhythm disruptions in the human brain associated with OUD, highlighting synaptic alterations in the dorsolateral prefrontal cortex (DLPFC) and nucleus accumbens (NAc)—key brain regions involved in cognition and reward, and heavily implicated in the pathophysiology of OUD. To provide further insights into the synaptic alterations in OUD, we used mass-spectrometry based proteomics to deeply profile protein expression alterations in bulk tissue and synaptosome preparations from DLPFC and NAc of unaffected and OUD subjects. We identified 55 differentially expressed (DE) proteins in DLPFC homogenates, and 44 DE proteins in NAc homogenates, between unaffected and OUD subjects. In synaptosomes, we identified 161 and 56 DE proteins in DLPFC and NAc, respectively, of OUD subjects. By comparing homogenate and synaptosome protein expression, we identified proteins enriched specifically in synapses that were significantly altered in both DLPFC and NAc of OUD subjects. Across brain regions, synaptic protein alterations in OUD subjects were primarily identified in glutamate, GABA, and circadian rhythm signaling. Using time-of-death (TOD) analyses, where the TOD of each subject is used as a time-point across a 24- hour cycle, we were able to map circadian-related changes associated with OUD in synaptic proteomes related to vesicle-mediated transport and membrane trafficking in the NAc and platelet derived growth factor receptor beta signaling in DLPFC. Collectively, our findings lend further support for molecular rhythm disruptions in synaptic signaling in the human brain as a key factor in opioid addiction.

## Introduction

Rates of opioid use disorder (OUD) and deaths from opioid overdoses have continued to rise in the United States over the recent decade (2012-2022) [1]. Further understanding of the impact of long-term opioid use and opioid overdose on the human brain is important for considering new avenues of treatment and intervention. Investigating the molecular alterations in postmortem brains from subjects with OUD provide valuable insights into the neurobiological mechanisms and potential therapeutic targets for opioid addiction [2–8].

Recent work in postmortem brains of subjects with OUD from us [2, 3] and others [4, 6, 9, 10] have primarily focused on the dorsolateral prefrontal cortex (DLPFC) and nucleus accumbens (NAc), key brain regions involved in cognition and reward, which are significantly impacted in OUD [11–13]. An interplay between inflammation and synaptic remodeling associated with OUD in both DLPFC and NAc has been consistent findings across human postmortem brain studies [3, 9, 10]. Most recently, our work linked alterations in neuroinflammatory, dopaminergic, and GABAergic signaling in DLPFC and NAc of OUD subjects with significant disruptions in circadian rhythms of transcript expression [2]. In OUD, alterations in transcriptional rhythms were associated with sleep and circadian traits (*e.g.,* insomnia) [2], further supporting biological relationships between changes in sleep, circadian rhythms, and synaptic signaling in opioid addiction [14–22].

To date, much of the work has used transcriptomics approaches to investigate the molecular alterations in postmortem brains of subjects with OUD, with limited studies using proteomics [10]. Complementary to transcriptomics, proteomics can report on functionally relevant information in cell bodies and neuropils, including synapses and glial processes [23–25]. Recent advances have enabled reliable detection of numerous peptides in synaptosomes isolated from postmortem human brains [24]. Deep profiling of protein changes at the synaptic level may lead to functional insights into disease-related mechanisms associated with psychiatric disorders.

Using quantitative mass spectrometry (MS) and tandem mass tags (TMT) [26, 27], we profiled both tissue homogenates and synaptosomes isolated from postmortem DLPFC and NAc of unaffected subjects to compare tissue-and synapse-level protein expression to subjects with OUD. Our approach allowed us to identify proteins that were preferentially expressed, or enriched, specifically in synapses of DLPFC and NAc, and assess synapse-specific alterations associated with OUD. Overall, pathways related to neuroinflammation, and neurodegeneration were altered across brain regions in OUD, accompanied by significant changes in proteins involved in GABAergic and glutamatergic synaptic signaling. Notably, synaptic alterations associated with OUD occurred in parallel to disruptions in pathways of circadian rhythm regulation. To further explore circadian rhythms in human postmortem brain, we used time-of-death (TOD) analysis to capture alterations associated with OUD in the diurnal variation of protein expression in synaptosomes. In synapses, disruptions to molecular rhythms were primarily related to endoplasmic reticulum functions, including protein trafficking and vesicle-mediated transport, platelet derived growth factor receptor (PDGFR) signaling, and proteins involved in GABA and glutamate synaptic signaling. Collectively, our results suggest protein pathways related to neurodegeneration are altered at the tissue level in DLPFC and NAc of OUD subjects, with implications for opioid-induced changes in circadian regulation of inhibitory and excitatory synaptic signaling.

## Materials and Methods

### Human Subjects

Following consent from next-of-kin, postmortem brains were obtained during autopsies at Allegheny County (Pittsburgh, PA, USA; N=39) or Davidson Country (Nashville, TN, USA; N=1) Medical Examiner’s Office. An independent committee of clinicians made consensus, lifetime DSM-IV diagnoses for each subject based on results from psychological autopsy, including structured family interviews, medical record reviews, and toxicological and neuropathological reports [28]. Similar procedures were used to confirm absence of lifetime psychiatric and neurological disorders in unaffected subjects. Procedures were approved by University of Pittsburgh Committee for Oversight of Research and Clinical Training Involving Decedents and Institutional Review Board for Biomedical Research. Cause of death in 19/20 OUD subjects was accidental, due to combined drug or opioid overdose. Accordingly, toxicology revealed that 19/20 OUD subjects had detectable opioids in their blood at time of death (Supplementary Table 1). Each OUD subject was matched with an unaffected comparison subject for sex, age, and postmortem interval (PMI) [2, 3]. Cause of death for unaffected subjects was either natural, accidental, or undetermined (Supplementary Table 1; opioids absent by blood toxicology). Cohorts differed by race (p=0.02) and brain pH (p=0.015, 0.2 pH units mean difference), and did not significantly differ in PMI, RNA integrity number (RIN), or TOD (p>0.25; Supplementary Table S1). TOD was determined from the death investigation report (Medical Examiner’s Office). DLPFC (Brodmann Area 9) and NAc were anatomically identified and collected, as previously described [2, 3].

### Brain Sample Preparation for Mass Spectrometry

Gray matter tissue (∼20mg) from DLPFC and NAc were collected from fresh-frozen coronal tissue blocks via cryostat to minimize contamination from white matter and other subregions [29, 30]. Homogenate and synaptosome preparations were obtained using a variation of our enrichment protocol for postmortem human brain tissues [24, 31, 32] with SynPER reagent (ThermoFisher). From each sample, 10µg total protein (as measured by Micro BCA) was reduced, alkylated, and trypsin digested on S-Trap™ micro spin columns (ProtiFi). Subject pairs were randomly assigned to TMT blocks and labeled with TMTPro channels 1-10, with brain regions and preparations assigned to separate blocks [33]. Additional aliquots from each sample were used for a pooled control, digested separately with S-Trap Midi™ columns, divided then labeled with TMTPro channels 1 and 12. TMT labeled preparations from the same block were pooled with 10µg of the labeled pooled controls. The TMT labeled peptide pools were separated into eight fractions with the Pierce™ High pH Reversed-Phase Peptide Fractionation Kit (ThermoFisher Scientific), evaporated, and reconstituted in 20 µl 97% H2O, 3% ACN, 0.1% formic acid.

### Mass Spectrometry

TMT labeled peptides (∼1µg) were loaded onto a heated PepMap RSLC C18 column (2□µm, 100 angstrom, 75□µm□×□50□cm; ThermoScientific), then eluted by gradients optimized for each high pH reverse-phase fraction [27]. Sample eluate was electrosprayed (2000□V) into an Orbitrap Eclipse Mass Spectrometer (MS; ThermoFisher Scientific) for analysis. MS1 spectra were acquired at a resolving power of 120,000. MS2 spectra were acquired in the Ion Trap with CID (35%) in centroid mode. Real-time search (RTS) (max search time□=□34□s; max missed cleavages□=□1; Xcorr□=□1; dCn□=□0.1; ppm□=□5) was used to select ions for SPS for MS3. MS3 spectra were acquired in the Orbitrap with HCD (60%) with an isolation window□=□0.7m/z and a resolving power of 60,000, and a max injection time of 400ms.

### Data Processing

Raw MS files were processed in Proteome Discoverer (v. 2.5; ThermoFisher Scientific). MS spectra were searched against the *Homo sapiens* SwissProt database. SEQUEST search engine was used (enzyme=trypsin, maximum missed cleavage=2, minimum peptide length=6, precursor tolerance=10ppm). Static modifications include acetylation (N-term, +42.011 Da), Met-loss (N-term, -131.040 Da), Met-loss+Acetyl (N-temr, -89.030 Da), and TMT labeling (N-term and K, +229.163 Da). Dynamic modification, oxidation (M, +15.995 Da). PSMs were filtered by the Percolator node (maximum Delta Cn=0.05, FDR =0.01). Reporter ion quantification was based on intensity values with the following settings/filters: integration tolerance=20ppm, method=most confident centroid, co-isolation threshold=100, and SPS mass matches [%] threshold = 65. Peptide intensity values were normalized within and across TMT plex runs with Normalization Mode=Total Peptide Amount and Scaling Mode=On All Average in Proteome Discoverer. Peptides used to sum to protein measures were determined using “Unique + Razor”. Only proteins of high confidence were retained for analyses and protein values were log2 transformed prior to analysis. The mass spectrometry proteomics data have been deposited to the ProteomeXchange Consortium via the PRIDE partner repository [34] with the dataset identifier PXD041333 and 10.6019/PXD041333.

### Differential Expression of Proteins

Limma-Voom with covariate selection (TMT plex, sex, age, PMI, and peptide expression) was used to detect differentially expressed (DE) proteins between OUD and unaffected subjects in homogenate and synaptosome fractions [24]. Proteins were considered DE if both p≤0.05 (unadjusted) and log_2_ fold-change (logFC) were greater than or equal to ±0.26 (20% change in expression), as previously used [3, 35, 36]. Over-representation pathway analyses were completed using clusterProfiler [37] and ReactomePA [38] for DE proteins. Rank-rank hypergeometric overlap (RRHO) was used to detect overlap of proteins expression changes between unaffected and OUD subjects. To assess preferential enrichment of proteins in synapses, we compared protein expression between homogenate and synaptosomes within unaffected subjects and separately, within OUD subjects. Protein abundance in synaptosomes was compared to abundance in homogenates to identify proteins with significantly greater levels of expression in synaptosomes (Bonferroni-corrected p<0.05). Proteins that showed a difference of at least 1.25-fold in the synaptosome relative to homogenates were classified as synapse-enriched proteins. All other proteins, by default, were defined as non-enriched. Analysis of synaptosome differences between unaffected and OUD subjects were limited to these enriched synaptic proteins, as described previously [24].

### Diurnal Rhythmicity Analysis

For each subject, TOD was converted to Zeitgeber Time (ZT) by using the times of sunrise and sunset on the day of death (ZT0 is sunrise, negative ZTs reflect hours prior to sunrise). Sinusoidal curves were fitted using nonlinear least-squares regression with the coefficient of determination used as a proxy of goodness-of-fit (R^2^). Estimates of empirical p-values were determined using null distributions of R^2^ generated from 1,000 TOD-randomized expression datasets. Diurnal rhythms of protein expression in homogenates and synaptosomes were identified separately in unaffected and OUD subjects. Rhythmic proteins were then compared between unaffected and OUD subjects (p<0.05; Fisher’s exact test). A complementary analysis determined which proteins exhibited a significant change in rhythmicity between unaffected and OUD subjects (ΔR^2^ = R^2^ Unaffected - R^2^ OUD). Change in rhythmicity analysis was restricted to proteins that were significantly rhythmic in unaffected or OUD subjects. Proteins were identified as significantly less rhythmic in OUD versus unaffected subjects if ΔR^2^ > 0 or more rhythmic if ΔR^2^ < 0. Null distributions of ΔR^2^ were permuted 1,000 times, whereby the unaffected and OUD subjects were permuted independently to generate null distributions (R^2^ unaffected and R^2^ OUD). Significantly less or more rhythmic in OUD was calculated by comparing the ΔR^2^ to the null ΔR^2^ for unaffected and OUD subjects. Differences in phase, amplitude, and base (mesor of fitted curve) were also calculated and restricted to significantly rhythmic proteins from both unaffected and OUD subjects. Pathway enrichments of rhythmic proteins were completed by Metascape (Gene Ontology; Hallmark; Reactome; Canonical; metascape.org) [39].

Heatmaps represent the top rhythmic proteins in homogenates and synaptosomes for each brain region, whereby each row represents a protein, and each column represents a subject, ordered by ZT. Protein expression was Z-transformed and ordered by phase (ZT of peak expression). Heatmaps were generated for: 1) top 200 rhythmic proteins in unaffected subjects; 2) top 200 rhythmic proteins identified in unaffected subjects then plotted for OUD subjects to assess circadian proteomic alterations in OUD; 3) top rhythmic proteins in OUD subjects; 4) top 200 rhythmic proteins in OUD subjects then plotted for unaffected subjects.

### Weighted Gene Co-Expression Network Analysis

To identify co-expression protein modules, we used weighted gene co-expression network analysis (WGCNA) on protein expression from homogenates or synaptosomes within each brain region and group [2, 3, 40–42]. Module differential connectivity (MDC) analyses examined the impact of OUD on protein co-expression network modules. For each module, the degree of connection was calculated in both unaffected and OUD subjects. MDC was defined as the ratio of the protein co-expression connections between unaffected and OUD subjects. To assess significance of MDC, we permuted (1000 times) the samples and module members to generate null distribution. When a module was built on unaffected subjects, a significant MDC greater than 1 indicated a gain of connectivity of the modules in OUD, while a significant MDC less than 1 indicated a loss of connectivity. Modules were tested for enrichment of DE proteins and rhythmic proteins in unaffected and OUD. ARACne was used to identify hub proteins in each module [43]. Proteins with an adjacency value higher than the 90^th^ quantile were considered neighbors. Hub proteins were identified with the highest number of N-hob neighborhood nodes (NHNN) than the average. Hub proteins specific to unaffected and OUD subjects were considered disease-specific hub proteins. Hub gene networks were plotted using Cytoscape (3.9.1). We then identified enriched pathways of synapse specific protein co-expression networks using SynGo [44].

## Results

### Brain Region-specific Protein Alterations in Tissue Homogenates and Synaptosomes associated with OUD in NAc and DLPFC

Between unaffected and OUD subjects, we investigated protein expression differences in tissue homogenates of DLPFC and NAc and isolated synaptosomes from each brain region. We identified 43 DE (p<0.05 and logFC±0.26) proteins (14 upregulated and 29 downregulated; Fig. 1a, Supplementary Tables S2, S3) in the NAc of subjects with OUD. In DLPFC, we identified 55 DE proteins, with most being upregulated in OUD (46 upregulated and 9 downregulated; Fig. 1b, Supplementary Tables S4, S5). In synaptosomes, we found 56 DE proteins (17 upregulated and 39 downregulated) in NAc (Fig. 1e, Supplementary Table S7, S8) and 161 DE proteins (80 upregulated and 81 downregulated) in DLPFC (Fig. 1f, Supplementary Table S9, S10).

**Figure 1.**
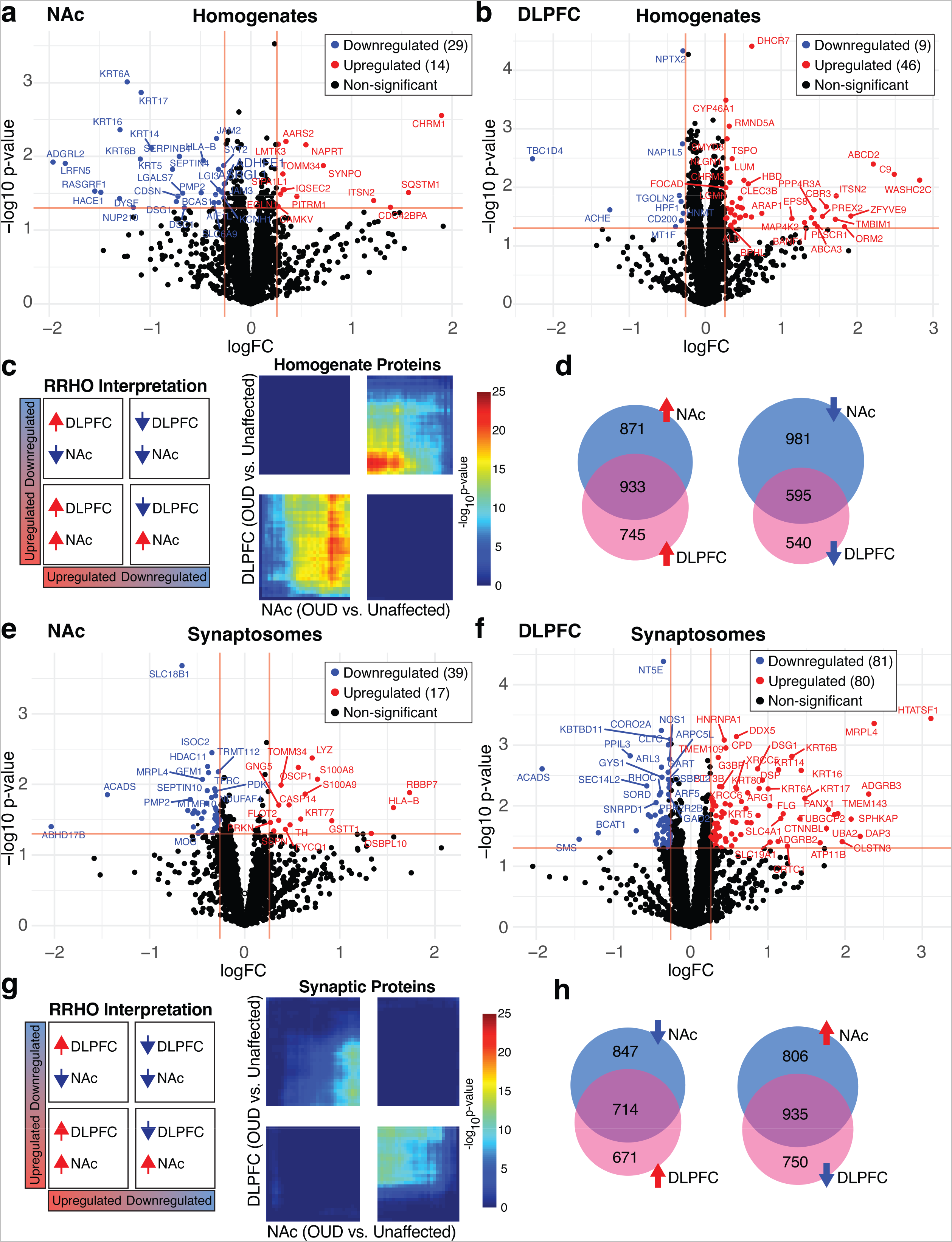
Proteomic alterations in tissue homogenates from NAc and DLPFC in subjects with OUD. **a**) Log_2_FC plotted relative to −log_10_ *p* value by volcano plot for DE proteins in NAc homogenates. **b**) Log_2_FC plotted relative to −log_10_ *p* value by volcano plot for DE proteins in DLPFC homogenates. **c**) RRHO plot indicating weak overlap or concordance of protein alterations between DLPFC and NAc in OUD subjects. **d**) Venn diagrams of downregulated and upregulated proteins between NAc and DLPFC homogenates. **e**) Log_2_FC plotted relative to −log_10_ *p* value by volcano plot for DE proteins in NAc synaptosomes. **f**) Log_2_FC plotted relative to −log_10_ *p* value by volcano plot for DE proteins in DLPFC synaptosomes. **g**) RRHO plot indicating very weak discordance of protein expression alterations in OUD between DLPFC and NAc synaptosomes. **h**) Venn diagrams of downregulated and upregulated proteins between NAc and DLPFC synaptosomes. Horizontal red lines indicate significance cutoffs of p<0.05, with vertical red lines represent log_2_FC±0.26 (panels a, b, e, and f). Proteins that reached both unadjusted p<0.05 and log_2_FC±0.26 were identified as DE proteins, upregulated labeled as red circles and downregulated labeled as blue circles.

To investigate the possible overlap of protein changes between brain regions, we compared protein expression between NAc and DLPFC of OUD subjects using the threshold-free approach, RRHO [45]. Overall, there was weak concordance of proteins that were upregulated or downregulated between brain regions of OUD subjects (Fig. 1c), despite identifying overlapping proteins between these brain regions (Fig. 1d). In contrast, we previously reported remarkable concordance between NAc and DLPFC in both upregulated and downregulated transcripts associated with OUD of the same subjects investigated here [2, 3]. In synaptosomes, we identified no significant overlap of protein changes between brain regions (Fig. 1g,h). Therefore, protein alterations in OUD were unique within brain region, and together with our previous results [3], may reflect the impact of opioids and other factors on brain region-specific, post-transcriptional and translational processing.

### Alterations in Inflammatory and Neurodegeneration-related Pathways associated with OUD in NAc and DLPFC Homogenates

To further define the biological significance of protein changes in OUD, we conducted pathway enrichment analysis on DE proteins from tissue homogenates of NAc and DLPFC (Fig. 2, Supplementary Table S6). In NAc homogenates, enrichment analyses identified pathways related to immune modulatory pathways, including various types of infection (*e.g.,* KRT14,16,17 [46]; HLA-B [47]) and leukocyte migration (*e.g.,* JAM2 [48]; PLCG1 [49]). Neurodegeneration-related pathways were also enriched in NAc homogenates (Fig. 2a; *e.g.,* CHRM1 [50, 51]; MAPT [52]; SQSTM1 [53]). Similarly, several pathways related to neurodegeneration were highly enriched in DLPFC homogenates from OUD subjects, such as Alzheimer (43 proteins; *e.g.,* SLC25A6 [54]; CHRM3 [55]; MTOR [56]; ITPR2 [57]), Huntington (38 proteins; *e.g.,* PSMD11 [58]; ADRM1 [59]; SOD1 [60]), Parkinson (41 proteins; PARK7 [61]; HTRA2 [62]), and prion (42 proteins; *e.g.,* RYR2 [63]) disease-related pathways (Fig. 2b). Other pathways enriched in DLPFC homogenates were related to cell stress including oxidative phosphorylation and reactive oxygen species (Fig. 2b). Together, our findings highlight a possible cascade of immune activation, diminished cellular health, and initiation of neurodegenerative processes in human brain that are associated with OUD.

**Figure 2.**
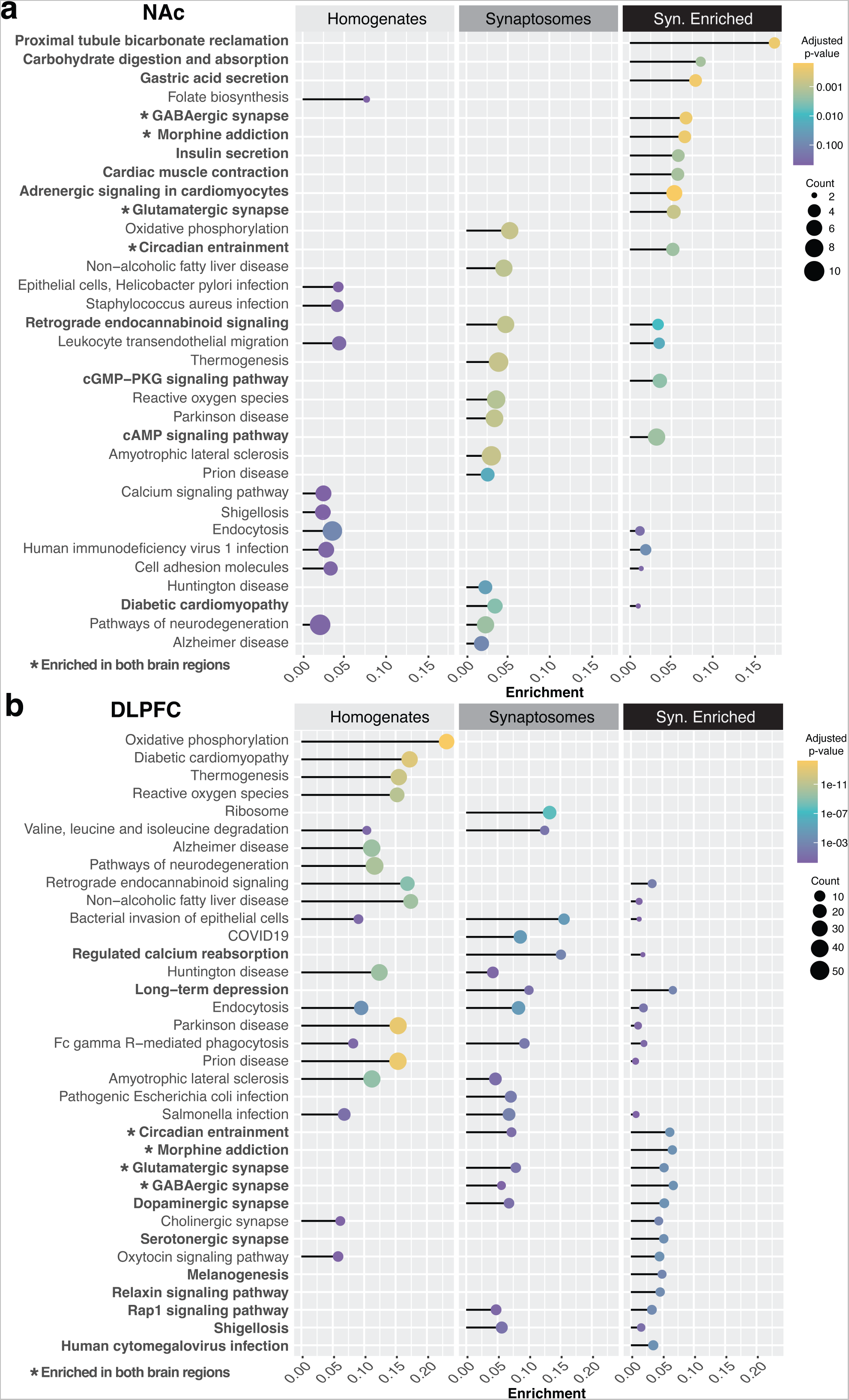
Differential enrichment of synaptic proteins between NAc and DLPFC associated with OUD. Lollipop plot showing pathways enriched from DE proteins in synaptosomes relative to homogenates in **a**) NAc and **b**) DLPFC. Adjusted p-value by color. Size of the circle represents counts of proteins within pathways. The enrichment score is calculated as the count of proteins identified as DE in OUD subjects divided by the count of proteins in the background of the respective ontological pathway. Bolded text with stars represents pathways enriched in both brain regions. Bolded only text represents pathways enriched in synaptosomes of the region analyzed. Also see Supplementary Table S11 for Enrichment in NAc Synaptosomes; Supplementary Table S12 for Enrichment in DLPFC Synaptosomes. NAc, Nucleus Accumbens; DLPFC, Dorsolateral Prefrontal Cortex.

### Differential Enrichment between Tissue Homogenates and Synaptosomes Highlights Altered GABAergic and Glutamatergic Synaptic Signaling in NAc and DLPFC associated with OUD

We investigated protein expression changes preferentially enriched in synaptosomes to identify synapse-specific proteome alterations associated with OUD. We identified proteins primarily expressed in homogenates and preferentially expressed in synaptosomes within each brain region (Supplementary Tables S7-S12). Synapse enriched proteins were then used for DE and pathway analyses comparing unaffected and OUD subjects. Synaptosomes isolated from NAc were enriched for several pathways involved in mitochondrial and endoplasmic reticulum functions (proximal tubule bicarbonate reclamation, carbohydrate digestion and absorption), along with second messenger signaling cascades (cGMP-PKG and cAMP: *e.g.,* GNAI2,3; GABRA2; ACY1; Fig. 2a, Supplementary Table S6). Other pathways were specifically enriched in DLPFC synaptosomes from OUD subjects, which included ribosome, relaxin, and Rap1 signaling (Fig. 2b). OUD-associated alterations in serotonergic synapses (*e.g.,* GNAO1; GNAS; GNAI2,3, GNB1,4) were also unique to DLPFC (Fig. 2b, Supplementary Table S6).

Additionally, we found similarly altered pathways in synaptosomes between brain regions. The major pathways enriched across NAc and DLPFC synaptosomes included morphine addiction and GABAergic signaling (*e.g.,* GNG3, GABBR2, GNB1 in NAc; and GNAO1, GNB4, SLC6A11 in DLPFC), and glutamatergic synaptic signaling (*e.g.,* GNG3, SLC1A3, and ADCY1 in NAc; and GNAO1, GNAS, and GNB14 in DLPFC; Fig. 2a,b).

Further investigation of the individual proteins enriched in synaptosomes revealed THY1 [64, 65], CACNA2D1 [66–70], SLC3A2 [70], GSK3B [71–73], LSAMP [73, 74], LY6H [75, 76], and NCAM1 [77–81], as the top synaptic proteins altered in NAc of OUD subjects (Fig. 3). In DLPFC, proteins altered in OUD and enriched in synaptosomes included RALB [82], CADM3 [83], GPC1 [84], MRAS, CNTFR [85], IGLON5, IST1, and PLXNA4 [86, 87] (Fig. 3). Other synapse enriched proteins in DLPFC were like those enriched in NAc (*e.g.,* GSK3B, LSAMP, NCAM1, CACNA2D1, SLC3A2; Fig. 3). Consistent with our pathway enrichments in synaptosomes of OUD subjects (Fig. 2), many of the synaptic proteins in both the NAc and DLPFC are involved in GABAergic and glutamatergic synaptic functions. For example, each of the synaptic proteins, CACNA2D1, SLC3A2, and GSK3B, altered in both brain regions, are known to dynamically regulate the activity of glutamatergic-dependent synaptic activity and plasticity [69, 70, 72].

**Figure 3.**
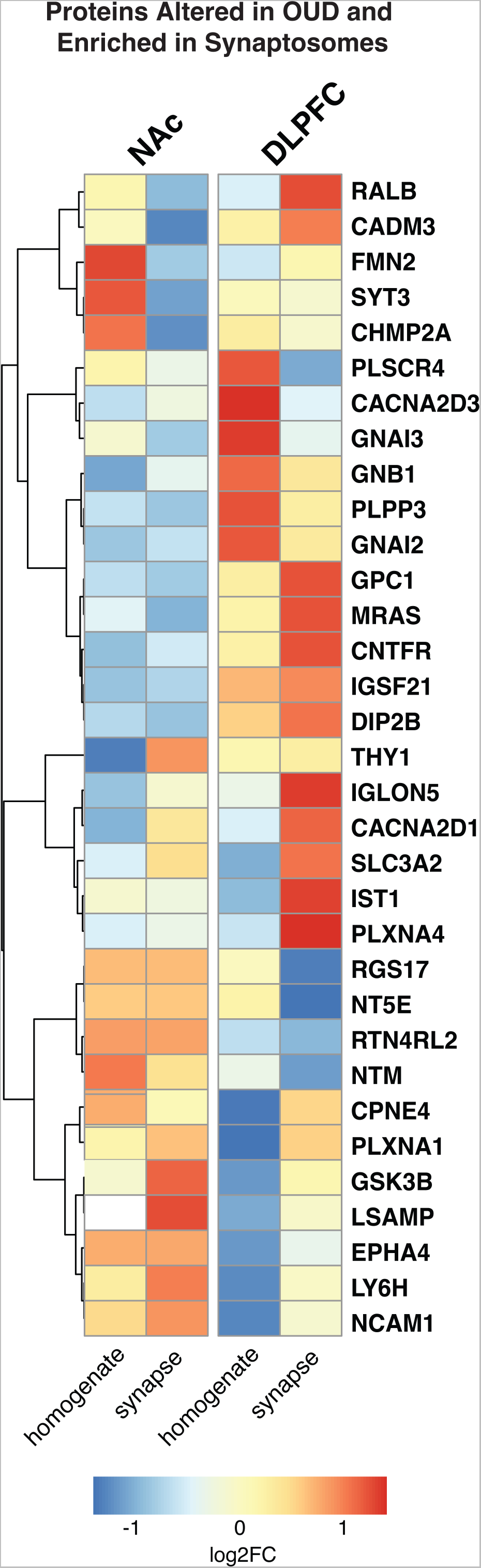
Alterations in protein synaptic enrichments in DLPFC and NAc associated with OUD. Heatmap highlighting top differentially expressed proteins in homogenates and synaptosomes in NAc and DLPFC of OUD subjects compared to unaffected controls. Warmer colors indicate increasing log_2_FC and highly enriched proteins in OUD. In contrast, cooler colors indicate decreasing log_2_FC and negative enrichment of proteins in OUD synaptosomes. Proteins are filtered for FDR < 0.10.

### Altered Circadian Rhythm Signaling associated with OUD in Synaptic Proteomes of NAc and DLPFC

Previously, we reported transcriptional rhythm disruptions in NAc and DLPFC of subjects with OUD [2]. In support of this, a significantly enriched pathway in DE synaptic proteins in both brain regions were circadian entrainment (Fig. 2). Several preferentially enriched synaptic proteins have integral roles in the regulation of circadian rhythms (*e.g.,* CACNA2D1, GSK3B, NCAM1, SYT3 [88]; Fig. 3). To further explore potential circadian rhythms in brain proteomes, we conducted TOD analysis on protein expression profiles from homogenate (Supplementary Fig. S2, Supplementary Fig. S1, Supplementary Tables S13-20) and synaptosomes (Fig. 4a and 4b) of each brain region. In synaptosomes, we identified 300 and 77 significantly rhythmic proteins in NAc of unaffected and OUD subjects, respectively (Fig. 4d). In DLPFC synaptosomes, we identified 123 rhythmic proteins in unaffected and 182 rhythm proteins in OUD (Fig. 4h). Top rhythmic synaptic proteins in unaffected subjects were significantly disrupted in subjects with OUD across both brain regions in synaptosomes (Figs. 4a,e; Supplementary Tables S17-S20 and S23-S24) and homogenates (Supplementary Fig S2; Supplementary Tables S13-S16 and S21-S22). Vice-versa, we also found that proteins were only rhythmic in OUD compared to unaffected subjects. Consequently, top rhythmic proteins in OUD were different from rhythmic proteins in unaffected subjects in synaptosomes (Fig 4a, NAc; Fig 4e, DLPFC; Supplementary Tables S17- S20 and S23-S24) and homogenates (Supplementary Fig. S1; Supplementary Tables S13-S16 and S21-S22). These findings highlighted a gain in rhythmicity in OUD compared to unaffected subjects.

**Figure 4.**
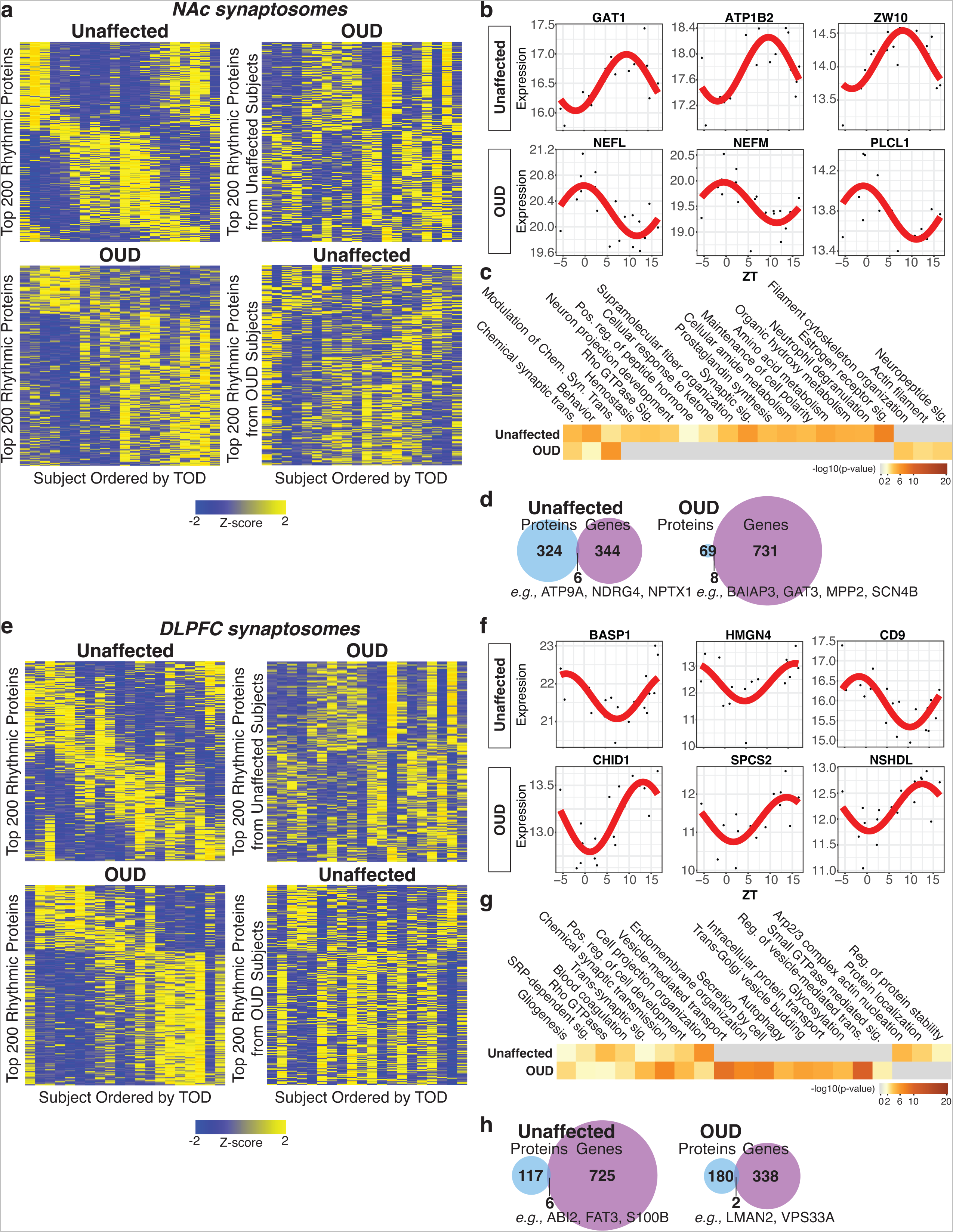
Diurnal rhythms of protein expression in DLPFC and NAc synaptosomes associated with OUD. **a**) Top left: Heatmap highlighting top 200 rhythmic proteins from NAc synaptosomes of unaffected subjects. Top right: Rhythmic proteins in unaffected subjects were plotted in OUD subjects to show disruption of protein rhythmicity. Bottom left: Heatmap highlighting top 200 rhythmic proteins from NAc synaptosomes of OUD subjects. Bottom right: rhythmic proteins in OUD subjects were plotted in unaffected subjects to show gain of protein rhythmicity in OUD. Heatmaps were generated by performing supervised clustering of expression of selected top 200 rhythmic proteins. Subjects were ordered by TOD to visualize expression levels over a period of 24 hours. Yellow color indicates increased Z-score and higher protein expression, while blue color indicates decreased Z-score and lower protein expression. **b**) Scatterplots of top rhythmic proteins in NAc synaptosomes from unaffected and OUD subjects. Scatterplots were generated to represent expression rhythms for individual proteins. The x-axis represents TOD on the ZT scale and protein expression level is on y-axis, with each dot representing a subject. The red line is the fitted sinusoidal curve to reflect temporal rhythms. **c**) Pathway enrichment analysis comparing rhythmic proteins in NAc synaptosomes from unaffected and OUD subjects. Warmer colors indicate increasing−log_10_ *p* value and highly rhythmic pathways in each group. **d**) Venn diagrams showing low overlap of rhythmic proteins and genes in NAc synaptosomes from unaffected and OUD subjects. **e**) Top left: Heatmap highlighting top 200 rhythmic proteins from DLPFC synaptosomes of unaffected subjects. Top right: Rhythmic proteins in unaffected subjects were plotted in OUD subjects to show disruption of protein rhythmicity. Bottom left: heatmap highlighting top 200 rhythmic proteins from DLPFC synaptosomes of OUD subjects. Bottom right: rhythmic proteins in OUD subjects were plotted in unaffected subjects to show gain of protein rhythmicity in OUD**. f**) Scatterplots of top rhythmic proteins in DLPFC synaptosomes from unaffected and OUD subjects. **g**) Pathway enrichment analysis comparing top 200 rhythmic proteins found in DLPFC synaptosomes from unaffected and OUD subjects. **h**) Venn diagrams showing low overlap of rhythmic proteins and genes in DLPFC synaptosomes from unaffected and OUD subjects. Also see Supplementary Figure S1 for protein expression rhythm heatmaps in unaffected homogenates; Supplementary Figure S2 for protein expression heatmaps in OUD homogenates; Supplementary table S13 for NAc Homogenates Rhythms in Unaffected subjects; Supplementary table S14 for NAc Homogenates Rhythms in OUD subjects; Supplementary table S15 for DLPFC Homogenates Rhythms in Unaffected subjects; Supplementary table S16 for DLPFC Homogenates Rhythms in OUD subjects; Supplementary table S17 for NAc Synaptosomes Rhythms in Unaffected subjects; Supplementary table S18 for NAc Synaptosomes Rhythms in OUD subjects; Supplementary table S19 for DLPFC Synaptosomes Rhythms in Unaffected subjects; Supplementary table S20 for DLPFC Synaptosomes Rhythms in OUD subjects; TOD, Time of Death; NAc, Nucleus Accumbens; DLPFC, Dorsolateral Prefrontal Cortex.

In NAc synaptosomes, the top rhythmic proteins of unaffected subjects included the GABA transporter, GAT1 [89, 90] (Fig. 4b). In OUD, top rhythmic proteins included neurofilament proteins (NEFL, NEFM [91]; Fig. 4b), consistent with enrichment of actin and filament cytoskeleton pathways (Fig. 4c). Neuropeptide signaling pathways were also enriched among rhythmic proteins in NAc of OUD subjects (Fig. 4c), a pathway that involves upstream initiation by G protein-coupled receptor binding to opioid receptors and downstream activation of intracellular pathways. Proteins encoding several opioid peptides were rhythmic in synaptosomes of OUD subjects, including PDYN (prodynorphin) and PENK (proenkephalin) [92, 93], and other proteins involved in opioid receptor signaling (SCG5 [94] and PCSK1N [95]). SCG5 and PCSK1N are also secretory proteins involved in preventing the aggregation of proteins involved in neurodegenerative disorders.

In DLPFC synaptosomes, top rhythmic proteins of unaffected subjects included brain abundant membrane attached signal protein 1, BASP1 (Fig.4f). Moreover, CHID1 [96] and SPCS2 [97] were among the top rhythmic synaptic proteins in DLPFC of OUD subjects (Fig. 4e,f), both of which are linked to cytoskeletal pathology in neurodegenerative disorders. Enrichment analyses of rhythmic synaptic proteins in DLPFC in OUD subjects revealed pathways primarily associated with Golgi and endoplasmic reticulum processing, autophagy, glycosylation, and synaptic vesicle transport (Fig. 4g). Disrupted endoplasmic reticulum signaling, along with changes in the process of protein glycosylation, may reflect the impact of opioids on local protein translation and trafficking at postsynaptic sites, including dendritic spines, regulating synaptic plasticity [98].

In NAc homogenates of OUD subjects, top enriched pathways included opioid and synaptic signaling (Supplementary Fig. S2). DARRP32, a phosphoprotein critically involved in dopaminergic synaptic functions, was among the top rhythmic proteins in OUD (Supplementary Fig. S2). Rhythmic proteins specifically in NAc of OUD subjects were associated with pathways in various vesicle endocytotic processes (*e.g.,* endocytosis, synaptic vesicle cycle, and clathrin-mediated endocytosis; Supplementary Fig. S2). In DLPFC, top rhythmic proteins in OUD subjects included the neurosecretory protein, VGF [99], and the putative negative regulator of cannabinoid receptor 1 activity, CNRIP1 (Supplementary Fig. S2). Top rhythmic pathways in OUD were related to glial cell development, mitochondrion organization, and various components of lipid and pyruvate metabolism (Supplementary Fig. S2). Rhythmicity in transcript abundance [2] was more prevalent compared to rhythms in protein expression in NAc and DLPFC (Supplementary Fig. S2). Notably, we observed very little overlap between rhythmic transcripts and rhythmic synaptic proteins (Fig. 4d, NAc; Fig 4h, DLPFC) and homogenates (Supplementary Figure S2) in either brain region of unaffected or OUD subjects.

### Altered Diurnal Rhythms of the Synaptic Proteome in DLPFC and NAc associated with OUD

Rhythmic proteins were largely distinct between unaffected and OUD subjects across brain regions. Given this, we examined whether proteins lost or gained rhythms between unaffected and OUD subjects within each brain region and sample preparation (Homogenates: Supplementary Figs. S1, S2, S3 and Supplementary Tables S21, S22; Synaptosomes: NAc, Fig. 5a; DLPFC, Fig. 5d; Supplementary Tables S23, S24). In NAc homogenates, 5 proteins lost rhythms and 19 proteins gained rhythms, primarily involved in protein translation (Supplementary Fig. S3a, S3c, Supplementary Table S21). Only 10 proteins were significantly altered in rhythmic expression in DLPFC homogenates of OUD subjects, with 3 proteins losing rhythms and 7 proteins gaining rhythms (Supplementary Fig. S3b). These proteins were enriched for pathways involved in lipid homeostasis and neurotoxicity, potentially related to neurodegenerative processes (Supplementary Fig. 3d; Supplementary table S22).

**Figure 5.**
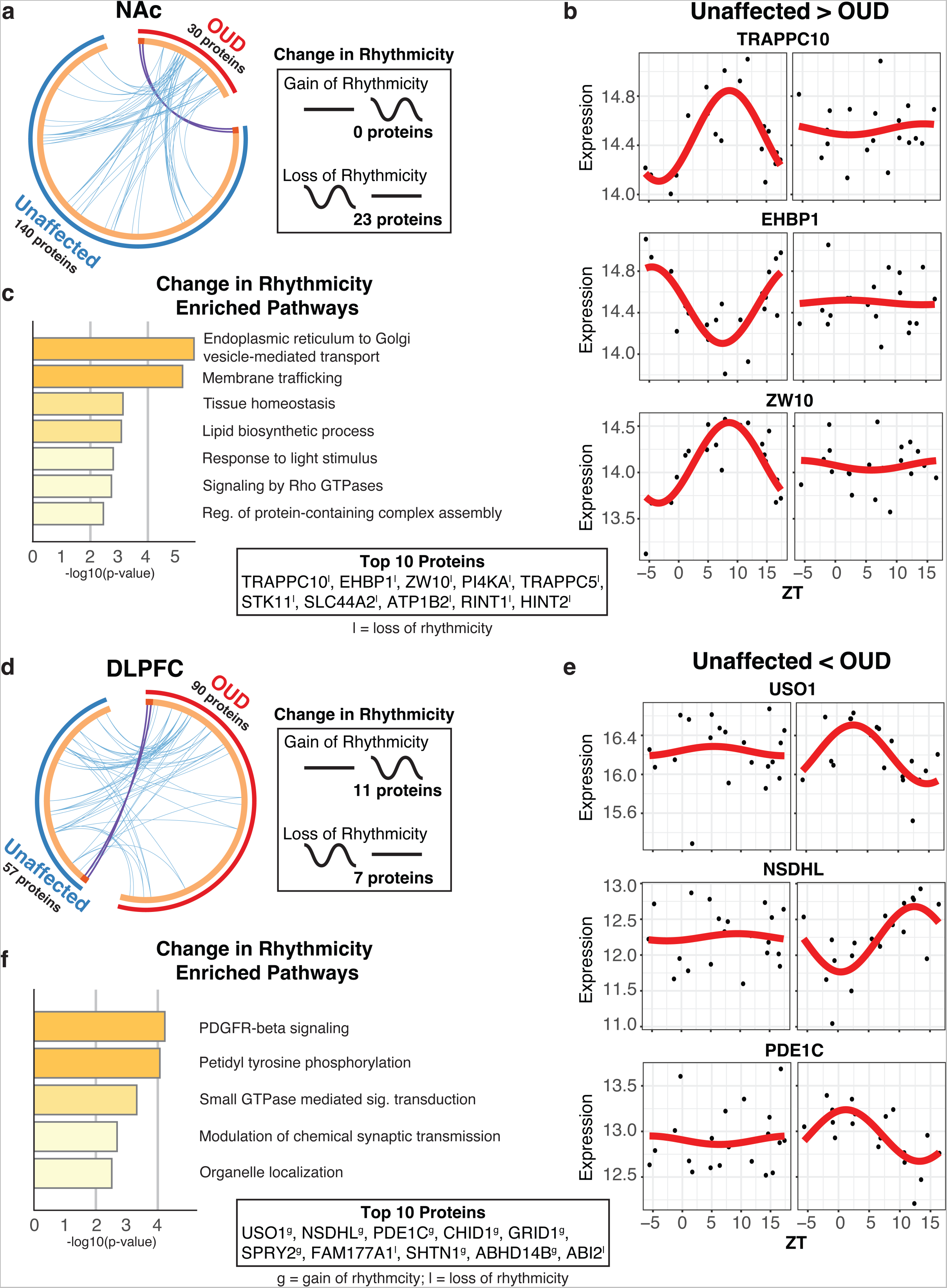
Altered rhythmicity of the synaptic proteome in DLPFC and NAc associated with OUD. Comparison of rhythmic proteins between unaffected and OUD subjects in synaptosomes from NAc and DLPFC. **a**) Circoplot highlights few rhythmic proteins that were identical (purple lines) and shared ontology (light blue lines) between unaffected and OUD subjects in NAc synaptosomes. Change in rhythmicity analysis in NAc Synaptosomes revealed that 0 proteins gained rhythmicity, while 23 lost rhythmicity in OUD subjects. **b**) Scatterplots of top proteins that lost rhythmicity in NAc synaptosomes between unaffected and OUD subjects. Scatterplots were generated to represent expression rhythms for individual proteins. The x-axis represents TOD on the ZT scale and protein expression level is on y-axis, with each dot representing a subject. The red line is the fitted sinusoidal curve to reflect temporal rhythms. **c**) Pathway enrichment on proteins that changed rhythmicity between unaffected and OUD subjects in NAc synaptosomes. **d**) Circoplot highlights few rhythmic proteins that were identical (purple lines) and shared ontology (light blue lines) between unaffected and OUD subjects in DLPFC synaptosomes. Change in rhythmicity analysis in DLPFC Synaptosomes revealed that 11 proteins gained rhythmicity, while 7 lost rhythmicity. **e**) Scatterplots of top proteins that lost rhythmicity in DLPFC synaptosomes between unaffected and OUD subjects. **f**) Pathway enrichment on proteins that changed rhythmicity between unaffected and OUD subjects in DLPFC synaptosomes. Also see Figure S3 for gain/loss rhythmicity proteins in NAc and DLPFC homogenates; Supplementary Figure S4 for DLPFC synaptosomes enriched gain and lost rhythm pathways; Table S21 for gain/loss rhythmicity protein list in NAc homogenates; Supplementary Table S22 for gain/loss rhythmicity protein list in DLPFC homogenates; Supplementary Table S23 for gain/loss rhythmicity protein list in NAc synaptosomes; Supplementary Table S24 for gain/loss rhythmicity protein list in DLPFC synaptosomes. TOD, time of death; NAc Nucleus Accumbens; DLPFC, DorsoLateral PreFrontal Cortex.

In synaptosomes, 23 proteins lost rhythmicity in NAc of OUD subjects (*e.g.,* TRAPPC10, EHBP1, ZW10; Fig. 5a,b; Supplemental Table S23). Loss of protein rhythms in NAc were enriched for endoplasmic reticulum to Golgi vesicle-mediated transport pathways (Fig. 5c). In DLPFC, we found 7 synaptic proteins that lost rhythmicity in OUD and 11 synaptic proteins that gained rhythmicity (Fig. 5d; *e.g.,* USO1, NSDHL, PDE1C [100], Fig. 5e). Proteins that gained rhythmicity (*e.g.,* USO1, NSDHL, PDE1C, CHID1, GRID1) were involved in chemical synaptic transmission and organelle localization, while proteins that lost rhythmicity (*e.g.,* FAM177A1, ABl2, TPD52L2, ElF2AK2, PRKCD) were involved in protein phosphorylation (Fig. 5f; Supplemental Table S24). Processed together, altered rhythmicity of synaptic proteins in DLPFC were associated with platelet-derived growth factor receptor beta (PDGFR-B) signaling (Fig. 5f), implicated in neuroprotection in response to elevated glutamatergic activity [101] and involved in opioid reward [102, 103]. Specifically, the 11 synaptic proteins gaining rhythmicity in DLPFC were associated with modulation of chemical synaptic transmission and organelle localization, while the 7 proteins losing rhythmicity were associated with protein phosphorylation (Supplementary Fig. S4; Supplemental Table S24). Collectively, our findings suggest circadian rhythm regulation of the synaptic proteome that is significantly altered in a brain region-specific manner in OUD. Further, circadian disruption of synaptic functions in OUD are associated with changes in endoplasmic reticulum-mediated local protein translation, trafficking, and vesicle endocytosis at the synapse, in addition to processes involved in neuroprotection and neurodegeneration.

### Protein Co-Expression Modules Identify Brain Region-Specific Alterations in Circadian Rhythm Signaling and Synaptic Functions associated with OUD

Using weighted co-expression analyses, we identified highly connected protein expression modules specific to NAc and DLPFC in homogenates (Supplementary Figs. S5, S7, S9; Supplementary Table S25) and synaptosomes (Supplementary Figs. S6, S8, S10; Supplementary Table S25). To further narrow modules relevant for OUD associated protein alterations, we used MDC analysis, which examined the overall connectivity of protein co-expression and network structure between unaffected and OUD subjects (Supplementary Table S26).

Many modules in unaffected subjects displayed a loss of connectivity in NAc and DLPFC homogenates (Supplementary Fig. S11; Supplementary Table S26) and synaptosomes (Fig. 6). In synaptosomes, connectivity was significantly lost in 13/15 modules in the NAc of OUD subjects (Fig. 6a), along with 10/26 modules in DLPFC (Fig. 6b). Loss of connectivity in protein co-expression modules indicates significant dispersion of protein network structure related to OUD.

**Figure 6.**
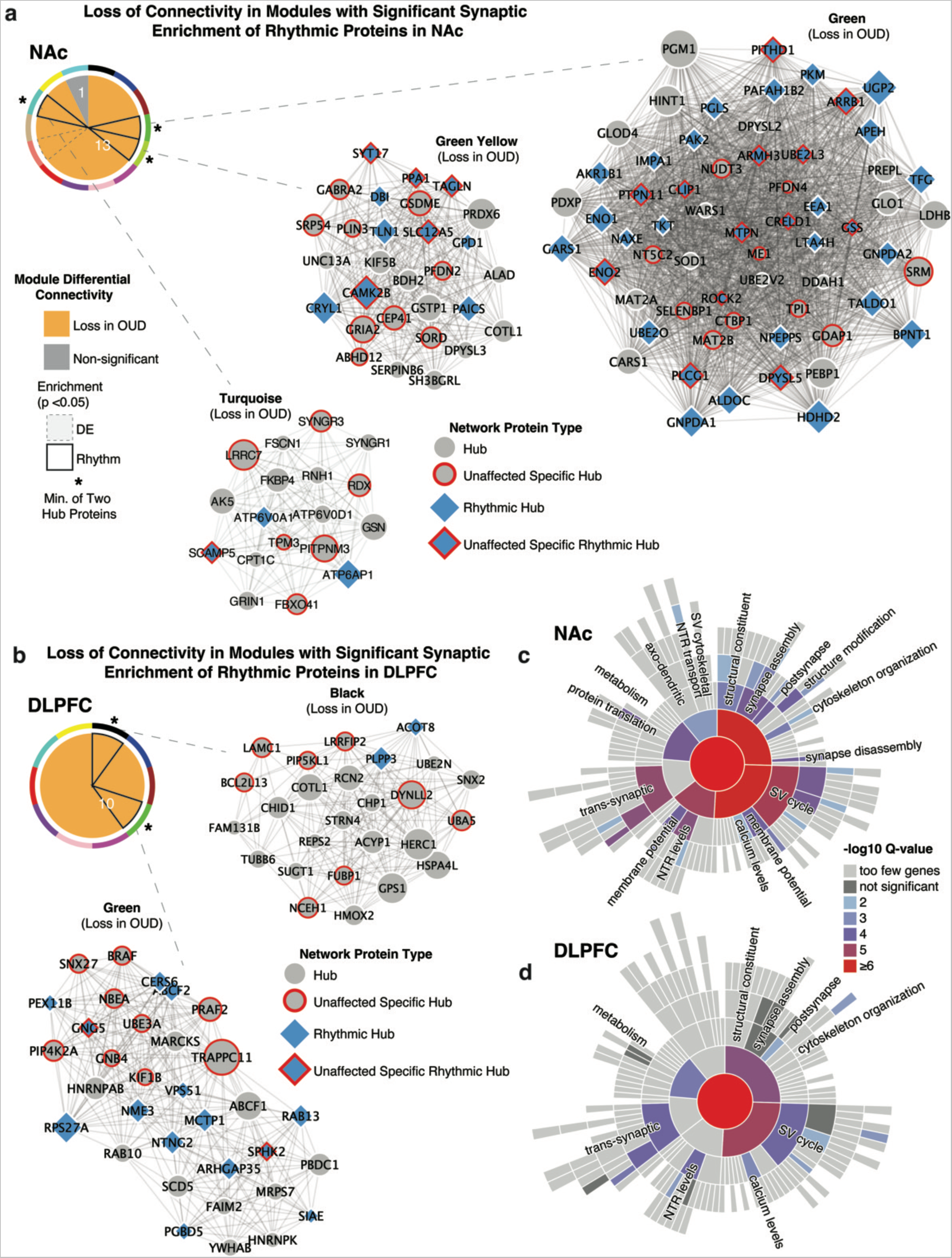
Altered protein networks in NAc and DLPFC synaptosomes associated with OUD. Weight Gene Correlation Network Analysis (WGCNA) was used to generate protein coexpression modules from each brain region separately. The identified modules that survived module preservation analysis were arbitrarily assigned colors. Pie charts generated from module differential connectivity (MDC) analysis summarize modules that gained or lost connectivity between unaffected and OUD subjects in NAc (**a**) and DLPFC (**b**). **a**) Comparing module connectivity between unaffected and OUD subjects in NAc synaptosomes identified 13 modules that lost connectivity in OUD, while 1 remained unchanged. Modules Turquoise, Green Yellow and Green were composed of several rhythmic protein hubs and showed loss in connectivity in OUD. **b**) Comparing module connectivity between unaffected and OUD subjects in DLPFC synaptosomes revealed all identified modules lost connectivity in OUD. Modules Green and Black were composed of several rhythmic hubs and showed loss of connectivity in OUD. **c**) Synaptic enrichment analysis of protein networks that lost connectivity in NAc synaptosomes. **d**) Synaptic enrichment analysis of protein networks that lost connectivity in DLPFC synaptosomes. Also see Figure S5 for WGCNA dendrograms on NAc and DLPFC homogenates; Figure S6 for WGCNA dendrograms on NAc and DLPFC synaptosomes; Figure S7 for modules from NAc homogenates; Figure S8 for modules from NAc synaptosomes; Figure S9 for modules from DLPFC homogenates; Figure S10 for modules from DLPFC synaptosomes; Figure S11 for additional MDC summaries; Supplementary Table S25 for WGCNA module assignments and proteins; Supplementary Table S26 for MDC analysis. DE, Differentially Expressed; Nucleus Accumbens; DLPFC, Dorsolateral Prefrontal Cortex.

We then investigated whether modules with changes in connectivity were enriched for hub proteins specific to unaffected and OUD subjects, and circadian rhythms. Focusing on the modules that lost connectivity in OUD, we found significant enrichment of both hub proteins specific to and/or rhythmic in unaffected subjects (Fig. 6). In NAc, there were three modules which lost connectivity in OUD that were significantly enriched for hub proteins (turquoise, green-yellow, and green; Fig. 6a). In the green module, proteins are associated with synaptic plasticity, dendritic spine formation, and axon guidance (*e.g.,* ROCK2 [104], DPYSL5 [105, 106], PLCG1 [107, 108], GDAP1; Fig. 6a). In the green-yellow module, several of the rhythmic hub proteins were related to glutamatergic (GRIA2 [109, 110]), GABAergic (GABRA2 [111–113]), and calcium signaling (CAMK2B, CALB2) (Fig. 6a). Additionally, other hub proteins are involved in calcium signaling, membrane potential, glutamatergic signaling, and exocytosis of synaptic vesicles (rhythmic hub SCAMP5; hubs SYNGR3 and LRRC7; Fig. 6a). Overall, protein networks that lost connectivity in the NAc of OUD subjects were enriched for hub proteins that exhibited rhythmicity in unaffected subjects and involved in GABAergic and glutamatergic neurotransmission (Fig. 6c).

In DLPFC, we identified two modules that lost connectivity in OUD and with significant hub protein enrichments (Fig. 6b). The black module contained several highly connected hub proteins involved in ECM formation (LAMC1 [114]), presynaptic dopamine release (PLPP3 [115]), neuroprotection (NCEH1 [116], BCL2CL13), inflammation (LRRFIP2 [117]), and pronociceptive receptor signaling in pain (PIP5KL1 [118]). The green module contained several rhythmic hub proteins involved neuronal protection (NME3, SPHK2, RAB13), synaptic functions (MCTP1, NTNG2, GNG5), and transposon activity in the brain (PGBD5 [119] (Fig. 6b). Enrichment analysis primarily highlighted synaptic functions of vesicle cycling and structure (Fig. 6d). Collectively, our analyses lend further support for a role of circadian rhythms in synaptic signaling within human brain and suggest significant disruptions of circadian-dependent regulation of synaptic functions in OUD.

## Discussion

Our previous transcriptomic findings implicate circadian rhythm disruption in proinflammatory signaling and synaptic remodeling in human NAc and DLPFC in OUD [2, 3]. We extend these findings here by showing significant alterations in circadian rhythm, GABAergic, and glutamatergic synaptic processes in both regions of OUD subjects. Using high-density profiling of peptides in tissue homogenates and synaptosomes within brain regions from the same subjects, we were able to identify and parse significant OUD-associated changes in synaptic proteomes specific to and common to NAc and DLPFC. In NAc synaptosomes specifically, pathways were involved in inflammatory, mitochondria, and metabolic signaling. In DLPFC synaptosomes from OUD subjects, pathways were mainly related to significant alterations in inflammatory signaling and serotonergic, dopaminergic, cholinergic, oxytocin neurotransmission. In both brain regions, pathways related to neurodegeneration, GABAergic and glutamatergic synapses, and circadian rhythms were associated with OUD, further supporting a role for circadian rhythms in synaptic signaling in opioid addiction.

Alterations in protein pathways related to neurodegeneration in OUD subjects were common in both NAc and DLPFC homogenates and synaptosomes. The relationship between chronic opioid use and increased risk for neurodegenerative disorders is complex. Several studies have reported increased risk of dementia and higher incidence of neurodegenerative disorders in people with heavy, chronic opioid use [120–122]. Chronic opioid use and dysfunction of the opioid system in the brain may contribute to neuroinflammation and the hyperphosphorylation of tau, a key protein involved in the pathogenesis of Alzheimer’s disease [122]. For example, dysfunction of opioid receptor signaling leads to increased amyloid beta expression and deposition [120]. Accumulation of amyloid beta is involved in the neurotoxicity observed early in the development of Alzheimer’s disease.

Consistent with the emergence of early pathogenesis associated with neurodegenerative disorders, several studies have shown significant elevation of early markers of neuropathology in Alzheimer’s disease in subjects who chronically used heroin, including hyperphosphorylated tau and amyloid beta [122]. Neurodegenerative markers in the brain were positively correlated with neuroinflammatory markers and microglial activation. Intriguingly, levels of the enzyme that phosphorylates tau, GSK-3B, were also increased in the same subjects [122]. We identified GSK-3B as a synaptic-enriched protein significantly upregulated in NAc and to a lesser degree in DLPFC of OUD subjects. Other related proteins were changed in both regions, including reduced expression of CBR1, BIN1, and VPS29 in DLPFC synaptosomes and reduced expression of SIRT2 in NAc synaptosomes. CBR1 is usually elevated in response to neuroinflammation and neuronal injury, ultimately facilitating neuroprotection [123]. Reduced BIN1 expression and function is a known risk factor for late-onset Alzheimer’s disease, having critical roles in presynaptic release of glutamate and the facilitation of learning and memory [124]. Further, VPS29 is involved in synaptic survival [125]. OUD-associated downregulation of each of these proteins in DLPFC may reflect the initiation of molecular signaling cascades resembling early pathogenesis of Alzheimer’s disease. In addition, SIRT2 downregulation in NAc synapses could lead to augmented activity at excitatory synapses via dysfunction of AMPA and NMDA glutamate receptors [126, 126, 127]. Accumulation of phosphorylated tau and other markers of neurodegenerative diseases, accompanied by elevated neuroinflammation in the brain, likely have functional impacts on synaptic processes in NAc and DLPFC in OUD. In OUD, several synaptic proteins were enriched in both NAc and DLPFC that link neurotoxicity to augmented glutamatergic neurotransmission–CNTFR, PLXNA4, and SLC3A2. Although speculative, alterations in synaptic processes we highlighted here in subjects with OUD, particularly GABAergic and glutamatergic neurotransmission, may be associated with cognitive, mood, and reward dysfunction associated with opioid addiction.

An imbalance of GABAergic and glutamatergic signaling in NAc and DLPFC has been linked to OUD and other psychiatric disorders. Between NAc and DLPFC of OUD subjects, our analyses identified several of the most impacted pathways in synaptosomes including GABAergic and glutamatergic neurotransmission. Several notable proteins involved in glutamatergic excitatory synapses were upregulated in DLPFC synaptosomes–GRIN2A (glutamate ionotropic receptor NMDA type subunit 2A), GRIN2B (glutamate ionotropic receptor NMDA type subunit 2A), GRM1 (glutamate metabotropic receptor 1), and SHANK2 (SH3 and multiple ankyrin repeat domains 2). GRIN proteins are subunits of the NMDA receptor complex, both of which are intimately involved in opioid-induced excitatory synaptic plasticity, having integral roles in opioid withdrawal and craving [128–130]. Synaptic elevation of SHANK2 may elevate NMDA receptor activity and enhance bursting of parvalbumin-positive neurons in DLPFC [131]. Enhancing NMDA receptor activation impairs synaptic plasticity and leads to cognitive impairments, potentially involved in OUD. We identified other synapse-specific proteins in the NAc involved in NMDA-dependent glutamate signaling, such as FLOT2 (upregulated), HDAC11 (downregulated), and SLC18B1 (downregulated). Notably, reduced HDAC11 expression in OUD and function may impede synaptic plasticity [132]. CACNA2D1, SLC3A2, and GSK3B, among others, were also enriched in both NAc and DLPFC of OUD subjects, each of which have dynamic roles in the regulation of glutamatergic-dependent synaptic plasticity. In addition, proteins involved in GABAergic processes were also significantly altered in OUD. For example, GAD2 was reduced in DLPFC synaptosomes in subjects with OUD. GAD2 is associated with synaptic vesicle response during intense neuronal activity to release GABA into the synaptic cleft [133], consistent with dysfunctional inhibitory neurotransmission in opioid addiction.

Additionally, we identified proteins specifically altered in NAc and DLPFC of OUD subjects. For example, LSAMP and LY6H were more enriched in NAc synaptosomes, both of which are involved in dendritic spine formation in response to excitatory inputs. Other proteins related to neuroinflammation were upregulated in NAc (*e.g.,* HLA-B, GSTT1, S100A9). In DLPFC, synaptic enriched proteins included IGLON5, a neuronal adhesion molecule linked to neurological disorder characterized by sleep disorders and cognitive impairments; EPHA4 [134], a receptor tyrosine kinase that modulates aberrant synaptic functions in response to neuronal injury and neuroinflammation; and GPC1, a glycoprotein involved in extracellular matrix (ECM) formation. Several proteins with the highest fold-change in OUD subjects included keratin cytoskeletal, ECM-related proteins in NAc [135]. Reduced expression and function of keratin proteins and neurofilaments are associated with neuropsychiatric disorders [121, 122], including stress-related disorders and substance use disorders [46]. The co-occurrence of changes in proteins involved in the activation of neuroinflammatory processes, degradation of ECM, and synaptic remodeling corroborates our previous transcriptomic findings in opioid addiction [3].

Pathway analysis implicated circadian rhythm disruption primarily in synaptosomes of OUD subjects. However, protein expression rhythms were markedly disrupted in both homogenates and synaptosomes from NAc and DLPFC. For example, DARPP-32, a protein highly expressed in striatal medium spiny neurons [136–138], was highly rhythmic in NAc homogenates, suggesting opioid-induced circadian reprogramming of molecular cascades in the striatum [139]. Notably, opioid signaling was a top pathway that was rhythmic in NAc homogenates, where rhythms in opioid neurotransmission may contribute to craving and relapse [140, 141]. Proteins that lost rhythmicity in the NAc of OUD subjects were involved in membrane trafficking and endoplasmic reticulum to Golgi vesicle-mediated transport (*e.g.,* TRAPPC5, TRAPPC10, EHBP1, ZW10). Both TRAPPC5 and TRAPPC10 are transmembrane proteins of the cis-Golgi complex that support vesicular transport from endoplasmic reticulum to Golgi. TRAPP10 interacts with TRAPPC9 to form TRAPP II core proteins. TRAPPC9/10 proteins regulate the endocytic receptor recycling of dopamine 1 and 2 receptors in postsynaptic striatal medium spiny neurons [142], integral to regulation of medium spiny neuron physiology and motivated behaviors. Local translation at synaptic sites may also occur through activity-dependent transport [143]. Glycoproteins THY1 and NCAM1 [144] were highly rhythmic in DLPFC homogenates. An area of future work could explore the impact of opioids on protein glycosylation of synaptic proteins and their impact of synaptic vesicle loading and release, particularly glutamate receptor turnover and activation [145] in response to neuroinflammation [146, 147]. In addition, PDGFR-B signaling was among the top pathways from altered rhythms in protein expression of DLPFC synaptosomes in OUD. Importantly, PDGF-dependent signaling is involved in numerous opioid actions, including tolerance and reward [148]. PDGF is a receptor tyrosine kinase, suggesting interactions between tyrosine kinases and opioid receptors may directly modulate glutamate receptor activity [149] depending on their circadian regulation.

Co-expression analyses revealed further insights into the role of circadian rhythm regulation of synaptic dysfunction in OUD. We identified several protein modules that were enriched for rhythmic proteins and hub proteins involved in synaptic processes in unaffected subjects (*e.g.,* CAMK2B, GSK-3B, ROCK2, PLCG1, SCAMP5, SYT17). Each of which are involved in dendritic spine formation and synaptic plasticity [104]. Every highly connected module in unaffected subjects in NAc and DLPFC lost connectivity in OUD subjects. Loss of connectivity in these protein networks in OUD could resemble a lack of coordinated protein signaling at the synapse that is modulated by circadian rhythms. OUD-associated upregulation of GSK-3B in synaptosomes of NAc and DLPFC likely leads to significant molecular rhythm disruptions in synapses, with potential consequences on activity-dependent synaptic plasticity, and thus, may be a lynchpin in opioid and circadian actions at the synapse.

Overall, our findings implicate circadian rhythm disruption in altered functions at GABAergic and glutamatergic synapses in both NAc and DLPFC. Considering the role of sleep and circadian rhythms in opioid reward, craving, and relapse, treatments that target circadian pathways may be an effective therapeutic strategy for OUD.

## Acknowledgments

Thank you to the staff and technicians who work diligently as part of the Brain Tissue Donation Program at the University of Pittsburgh. Postmortem human brain tissue was provided by the University of Pittsburgh Brain Tissue Donation Program and the National Institutes of Health NeuroBioBank at the University of Pittsburgh.

## Author Contributions

MLS and RWL obtained funding for the project. JRG, DAL, MLS, MLM, and RWL designed and coordinated the study. SP, RS, MAS, MAH, SMK, JRG, and MLM processed samples for proteomics. SP, XX, ZF, GCT, AY, MLM, and RWL conducted analyses and data interpretation. SP, XX, MLS, MLM, and RWL wrote the manuscript. All authors read and approved the final version of the manuscript.

