## Supplemental Material for "Circadian rhythm disruptions associated with opioid use disorder in the synaptic proteomes of the human dorsolateral prefrontal cortex and nucleus accumbens"

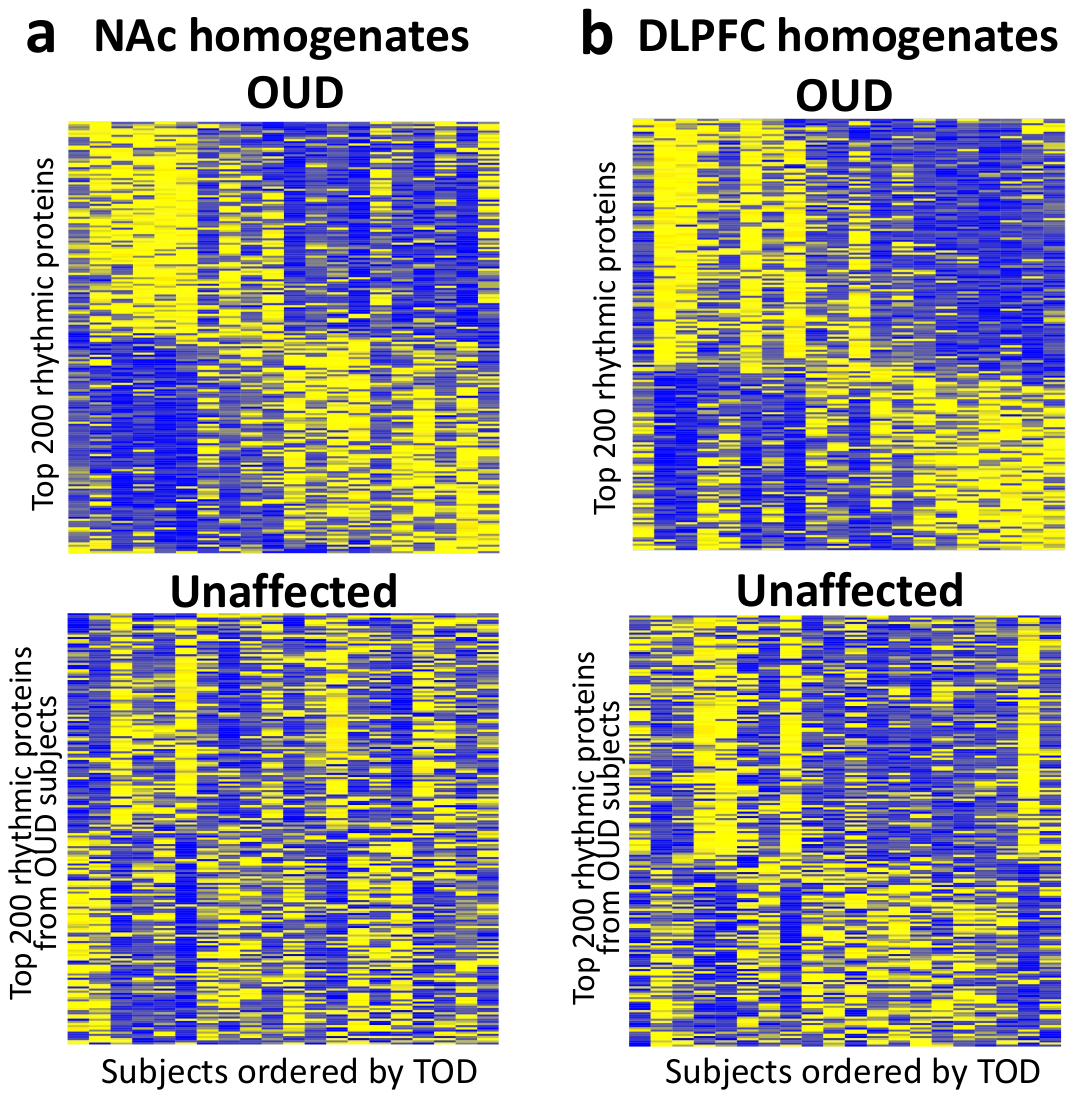

**Figure S1. Protein rhythm heatmaps.** **a**) Heatmaps highlighting the top 200 OUD rhythmic proteins found in NAc homogenates. **b**) Heatmaps highlighting that the top 200 OUD rhythmic proteins found in DLPFC homogenates. Heatmaps were generated by performing supervised clustering of expression of top 200 rhythmic proteins found in homogenates from unaffected subjects and applied to OUD. Subjects were ordered by TOD to visualize expression levels over 24-hours. Yellow color indicates increased Z-score and higher protein expression, while blue color indicates decreased Z-score and lower protein expression. TOD, time of death; NAc, nucleus accumbens; DLPFC, dorsolateral prefrontal cortex.

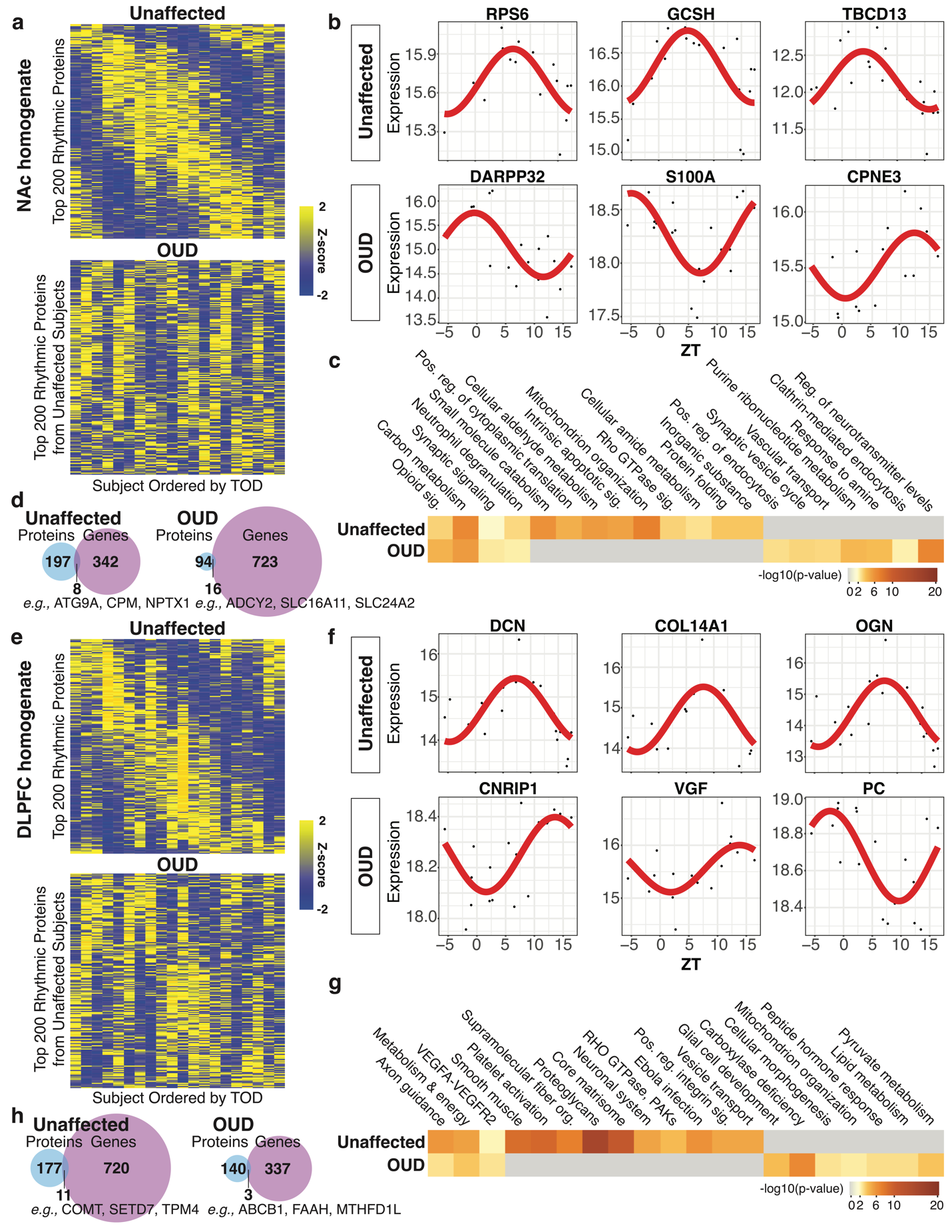
**Figure S2. Diurnal rhythms of protein expression in DLPFC and NAc homogenates associated with OUD.** **a**) Heatmaps highlighting top 200 rhythmic proteins from NAc homogenates of unaffected subjects. Rhythmic proteins in unaffected subjects are plotted in OUD subjects to show disruption of protein rhythmicity. Heatmaps were generated by performing supervised clustering of expression of top 200 rhythmic proteins found in homogenates from Unaffected subjects and applied to OUD. Subjects were ordered by TOD to visualize expression levels over a period of 24 hours. Yellow color indicates increased Z-score and higher protein expression, while blue color indicates decreased Z-score and lower protein expression. **b**) Scatterplots of top rhythmic proteins in NAc homogenates of unaffected and OUD subjects. Scatterplots were generated to represent expression rhythms for individual proteins. The x-axis represents TOD on the ZT scale and protein expression level is on y-axis, with each dot representing a subject. The red line is the fitted sinusoidal curve to reflect temporal rhythms. **c**) Pathway enrichment analysis of rhythmic proteins in NAc homogenates from unaffected and OUD subjects. Warmer colors indicate increasing −log_10_ *p* value and highly rhythmic pathways in each group. **d**) Venn diagrams showing low overlap of rhythmic proteins and genes in NAc homogenates from unaffected and OUD subjects. **e**) Heatmaps highlighting disrupted protein rhythmicity in OUD of top 200 rhythmic proteins from DLPFC homogenates of unaffected subjects. Rhythmic proteins in unaffected subjects are plotted in OUD subjects to show disruption of protein rhythmicity. **f**) Scatterplots of top rhythmic proteins in DLPFC homogenates from unaffected and OUD subjects. **g**) Pathway enrichment analysis of rhythmic proteins in DLPFC homogenates from unaffected and OUD subjects. **h**) Venn diagrams showing low overlap of rhythmic proteins in DLPFC homogenates from unaffected and OUD subjects. Also see Figure S1 for protein expression rhythm heatmaps in OUD; Supplementary Table S15 for rhythmic proteins in NAc homogenates of unaffected subjects; Supplementary Table S16 for rhythmic proteins in NAc homogenates of OUD subjects; Supplementary Table S17 for rhythmic proteins in DLPFC homogenates of unaffected subjects; Supplementary Table S18 for rhythmic proteins in DLPFC homogenates of OUD subjects. TOD, Time of Death; NAc, Nucleus Accumbens; DLPFC, Dorsolateral Prefrontal Cortex.

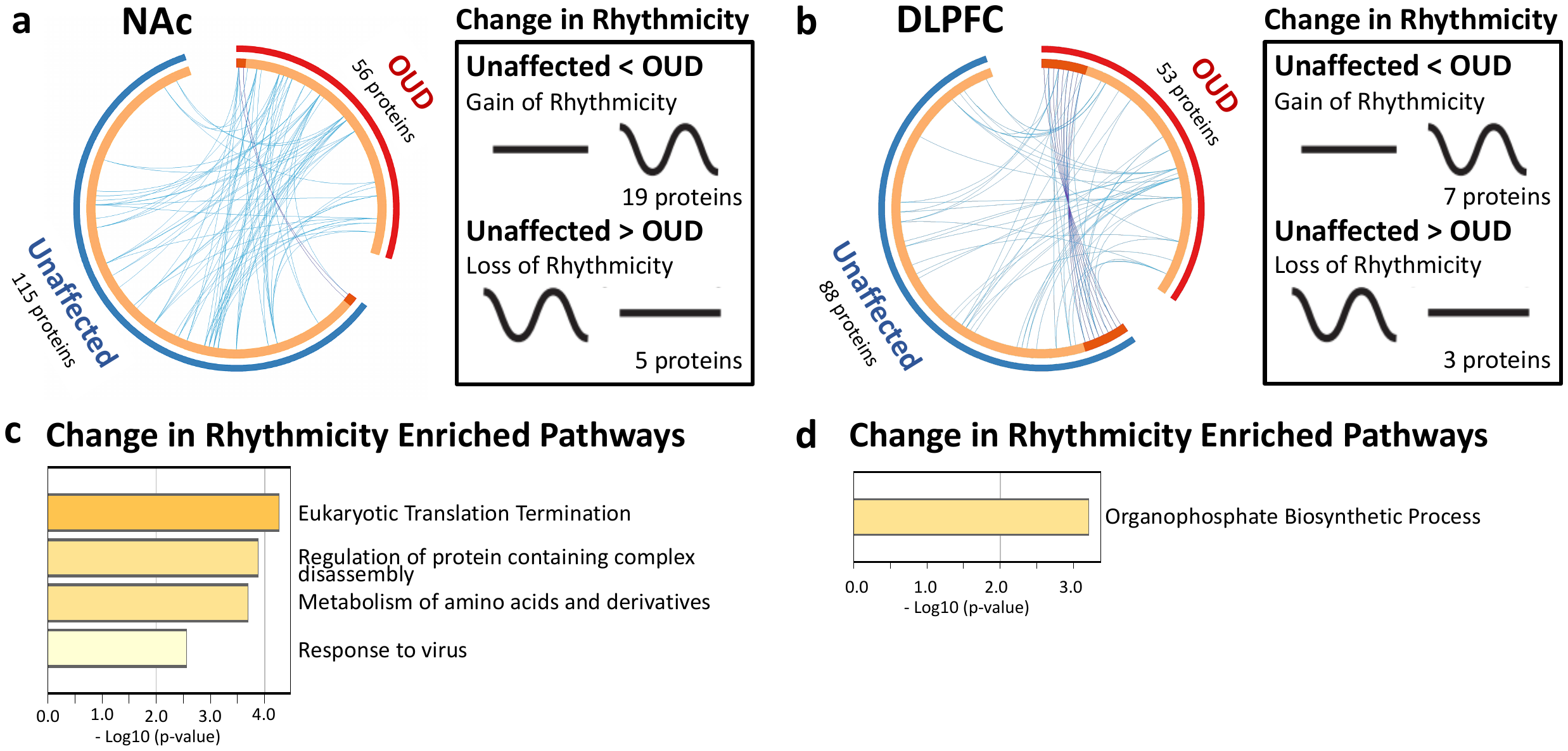

**Figure S3. Gain/loss of protein rhythms in NAc and DLPFC homogenates**. Comparison of rhythmic proteins from unaffected and OUD subjects in NAc and DLPFC homogenates. Circoplots highlight that few rhythmic proteins were identical (purple lines) and some shared ontology (light blue lines) between the DLPFC and NAc. **a**) Change in rhythmicity analysis in NAc homogenates revealed that 19 proteins gained rhythmicity, while 5 lost rhythmicity. **b**) Change in rhythmicity analysis in DLPFC homogenates revealed that 7 proteins gained rhythmicity, while 3 lost rhythmicity. **c**) Analysis of enriched pathways in proteins changing rhythmicity between unaffected and OUD in NAc homogenates. **d**) Analysis of enriched pathways in proteins changing rhythmicity between Unaffected and OUD in DLPFC homogenates.

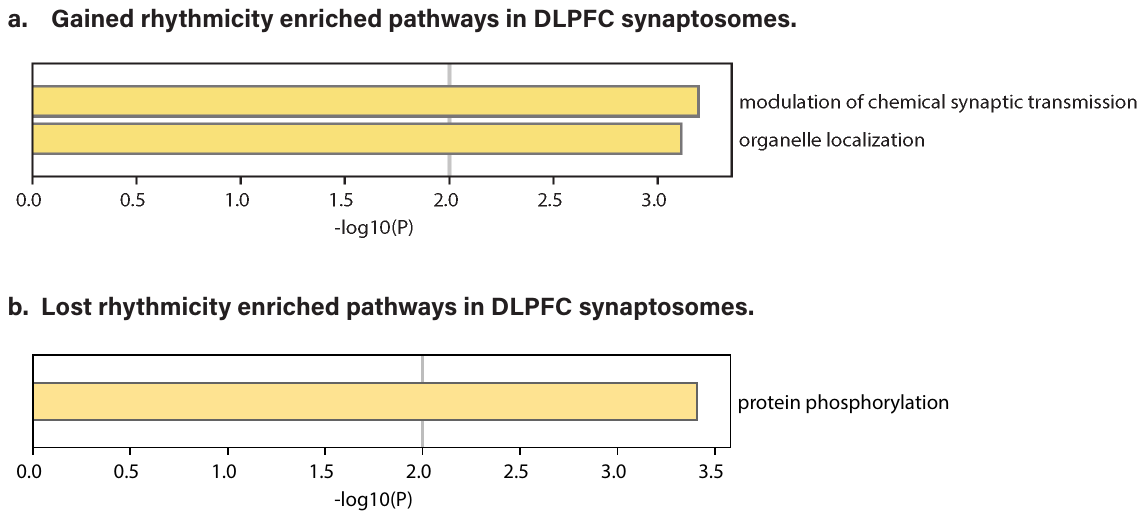

**Figure S4. Pathway enrichment on proteins with altered rhythmicity in DLPFC synaptosomes.**

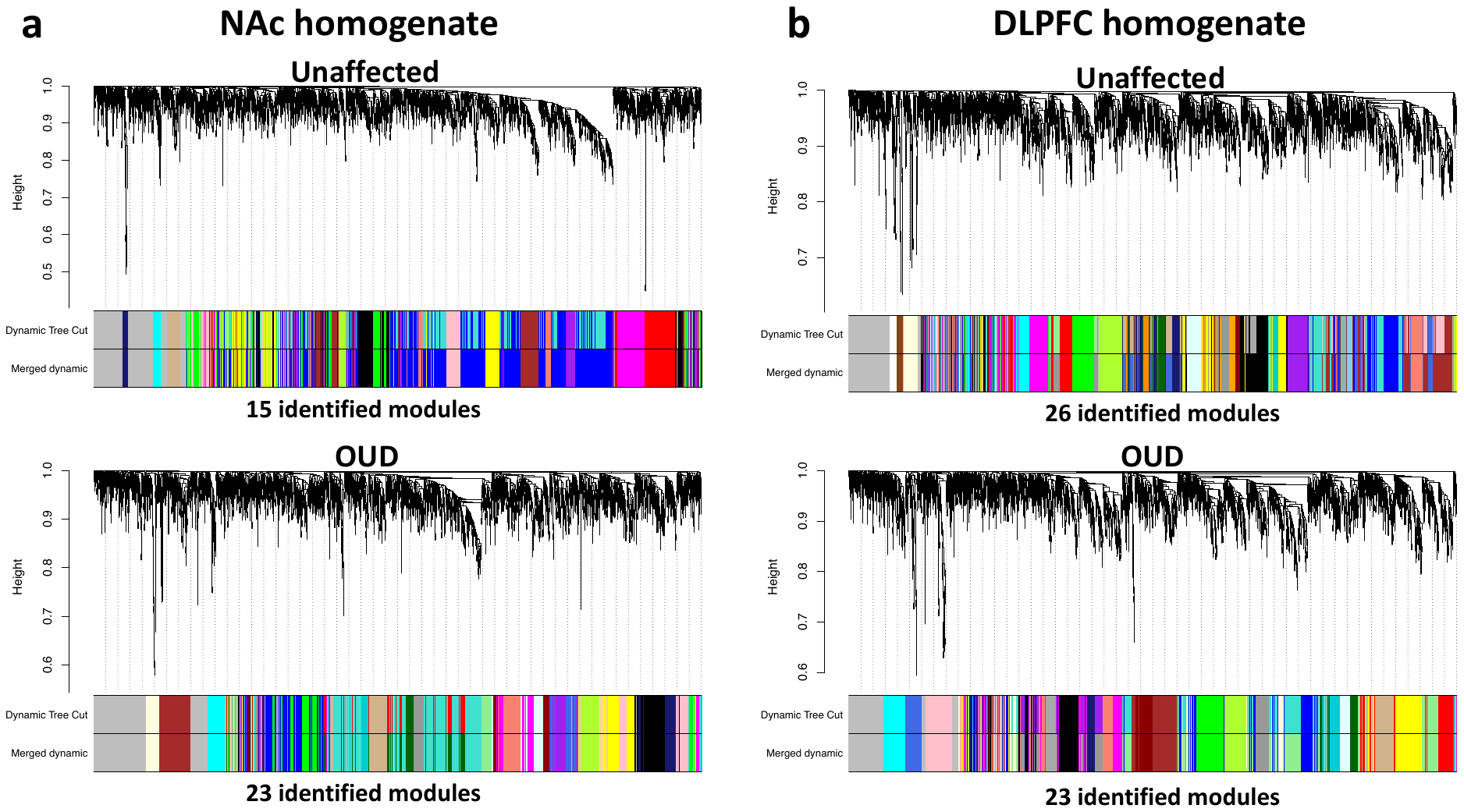

**Figure S5. Protein networks in NAc and DLPFC homogenates.** Dendrograms from WGCNA showing the average linkage hierarchical clustering of proteins. **a**) Dendrograms from NAc homogenates identified 15 and 23 modules in unaffected and OUD subjects, respectively. **b**) Dendrograms from DLPFC homogenates identified 26 and 23 modules in unaffected and OUD subjects, respectively. NAc, Nucleus Accumbens; DLPFC, Dorsolateral Prefrontal Cortex.

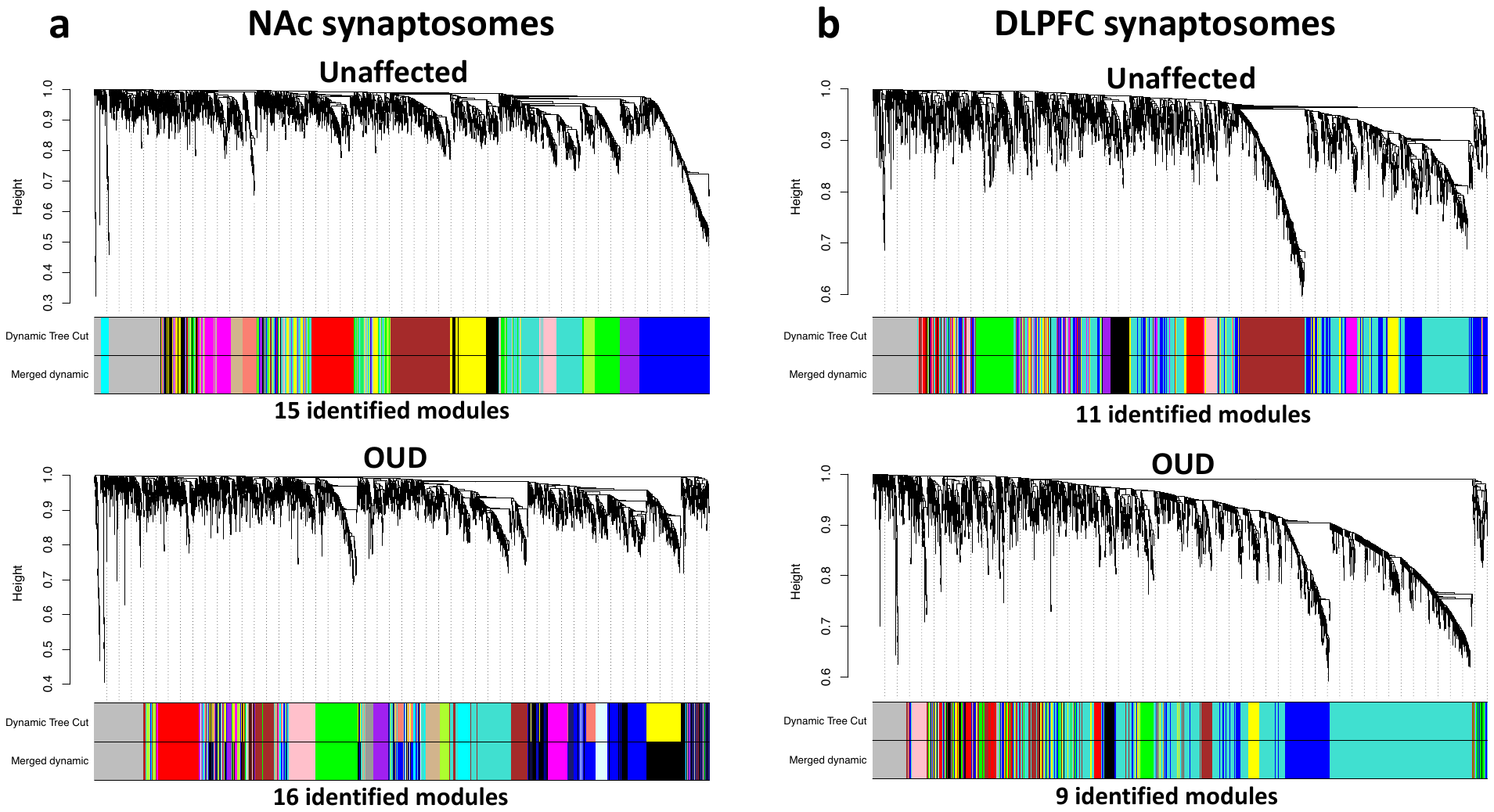

**Figure S6. Protein networks in NAc and DLPFC synaptosomes.** Dendrograms from WGCNA showing the average linkage hierarchical clustering of genes. **a**) Dendrograms from NAc Synaptosomes identified 15 and 16 modules in unaffected and OUD subjects, respectively. **b**) Dendrograms from DLPFC synaptosomes identified 11 and 09 modules in unaffected and OUD subjects, respectively. NAc, Nucleus Accumbens; DLPFC, Dorsolateral Prefrontal Cortex.

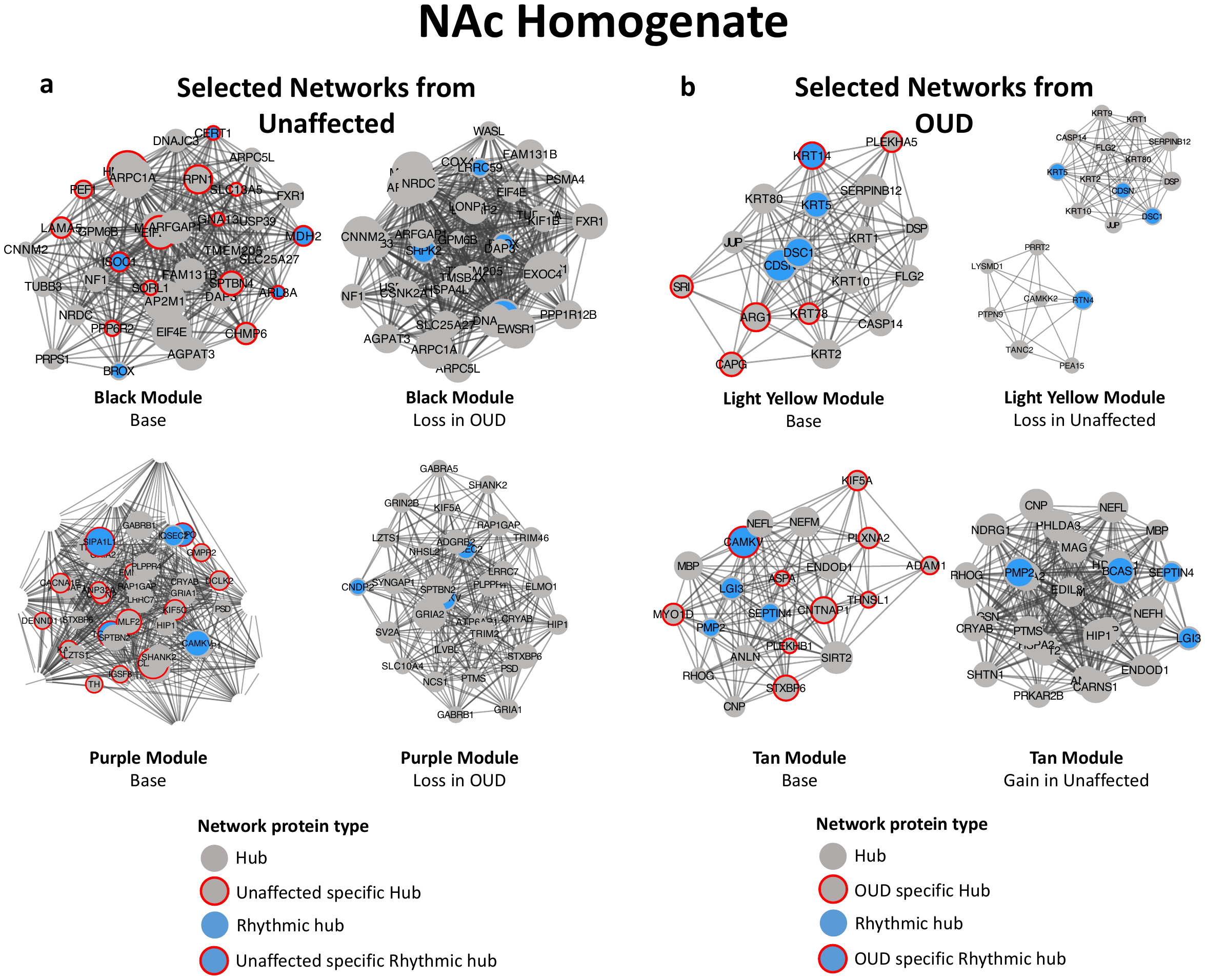

**Figure S7. NAc homogenate modules.** Selected networks from NAc homogenate modules of interest and significant hub enrichment of rhythmic proteins. **a**) Selected networks from base modules in unaffected to which were compared OUD. **b**) Selected networks from base modules OUD to which were compared unaffected.

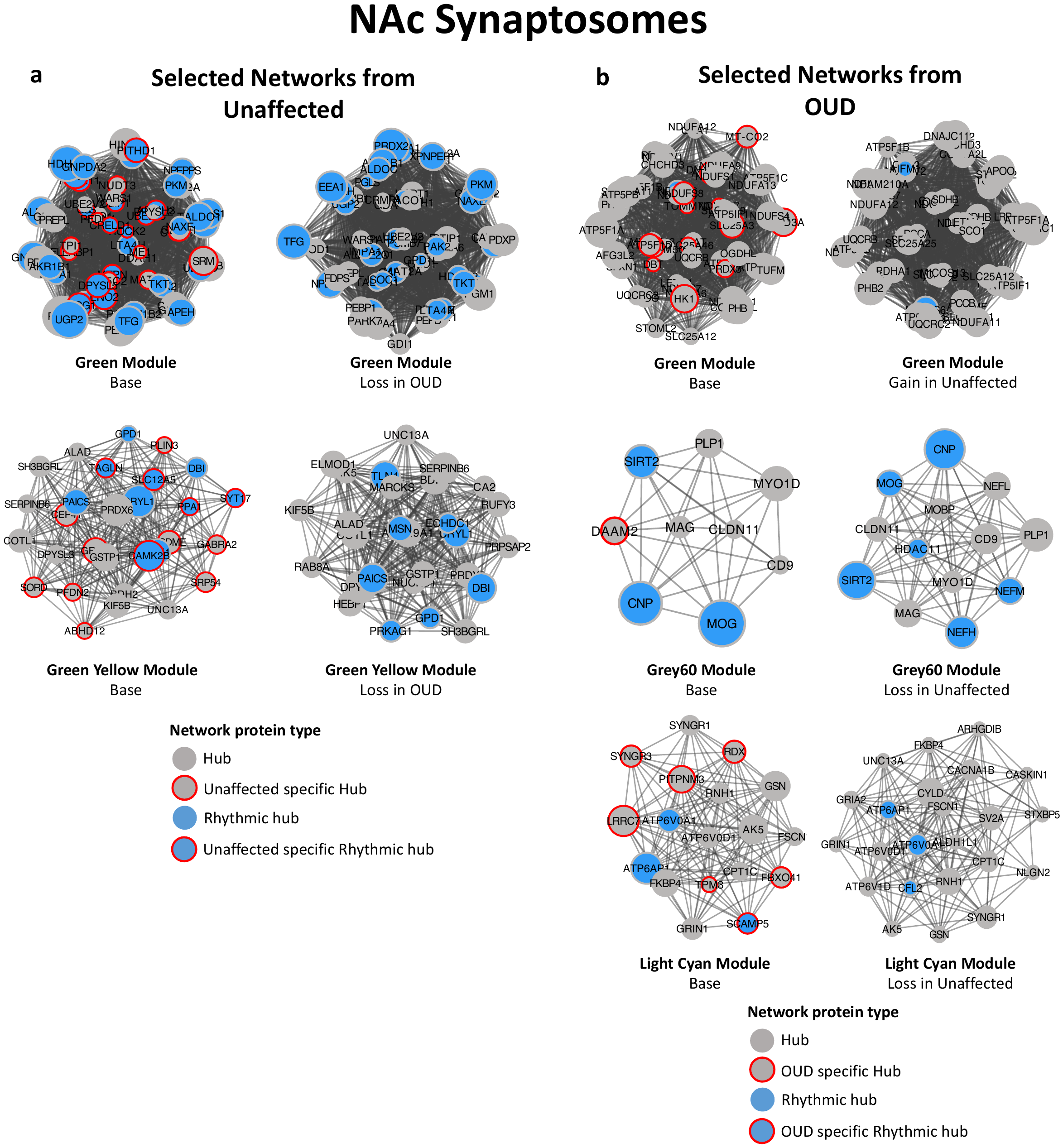

**Figure S8. NAc synaptosome modules.** Selected networks from NAc synaptosome modules of interest and significant hub enrichment of rhythmic proteins. **a**) Selected networks from base modules in unaffected to which were compared OUD. **b**) Selected networks from base modules OUD to which were compared unaffected.

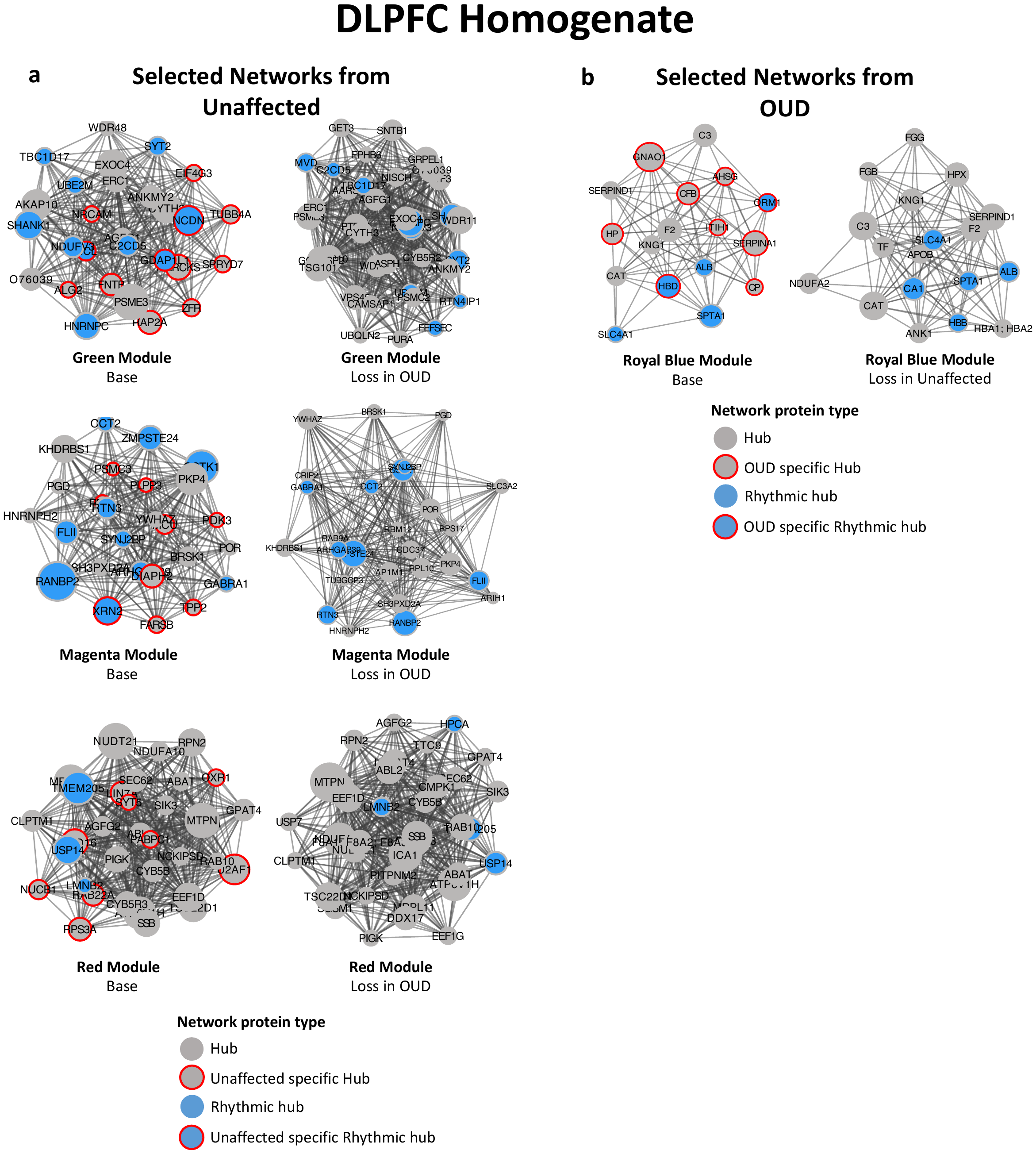

**Figure S9. DLPFC homogenate modules.** Selected networks from DLPFC homogenates modules of interest and significant hub enrichment of rhythmic proteins. **a**) Selected networks from base modules in unaffected to which were compared OUD. **b**) Selected networks from base modules OUD to which were compared unaffected.

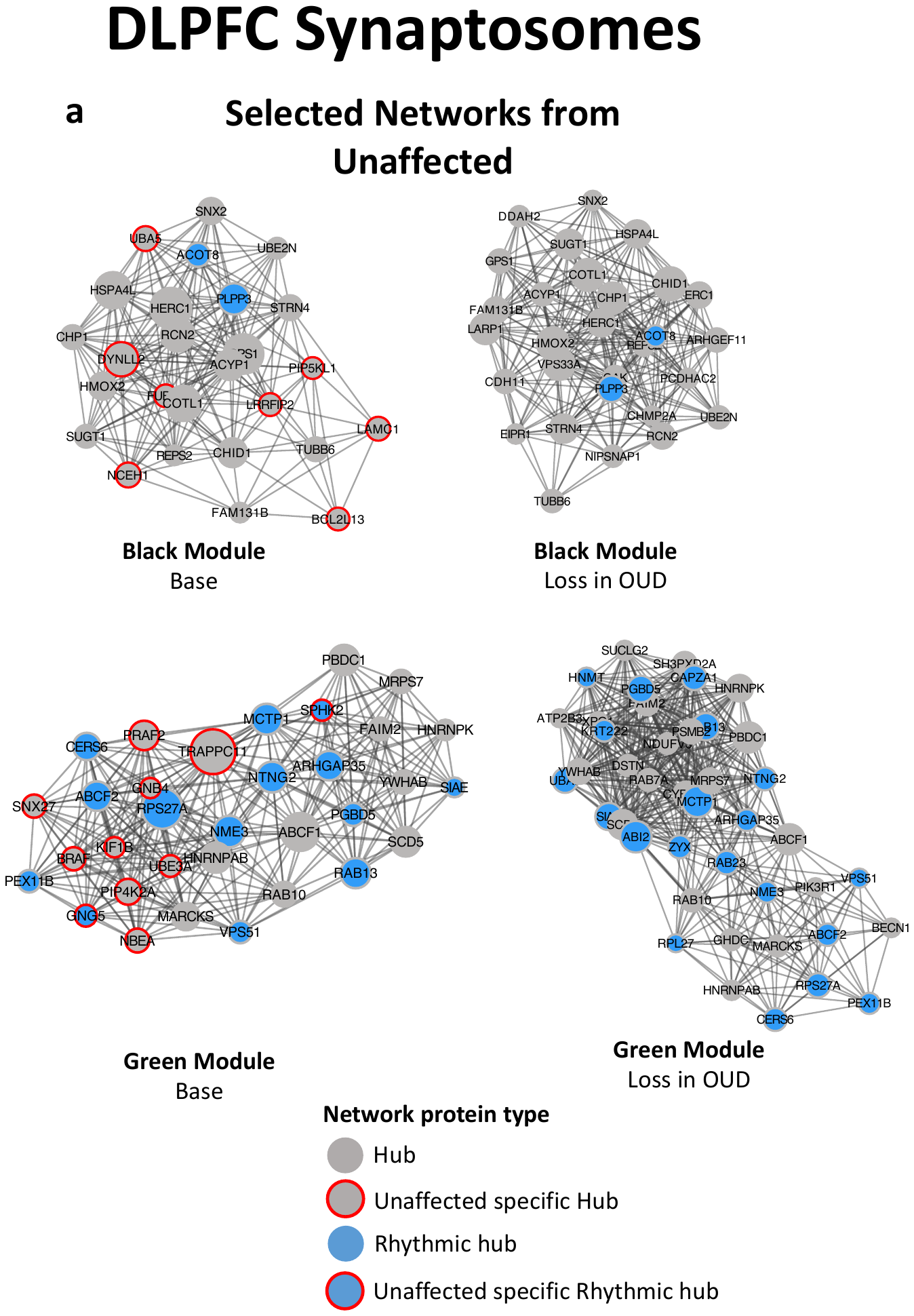

**Figure S10. DLPFC synaptosomes modules.** Selected networks from DLPFC synaptosomes modules of interest and significant hub enrichment of rhythmic proteins. **a**) Selected networks from base modules in unaffected to which were compared OUD. **b**) Selected networks from base modules OUD to which were compared to unaffected.

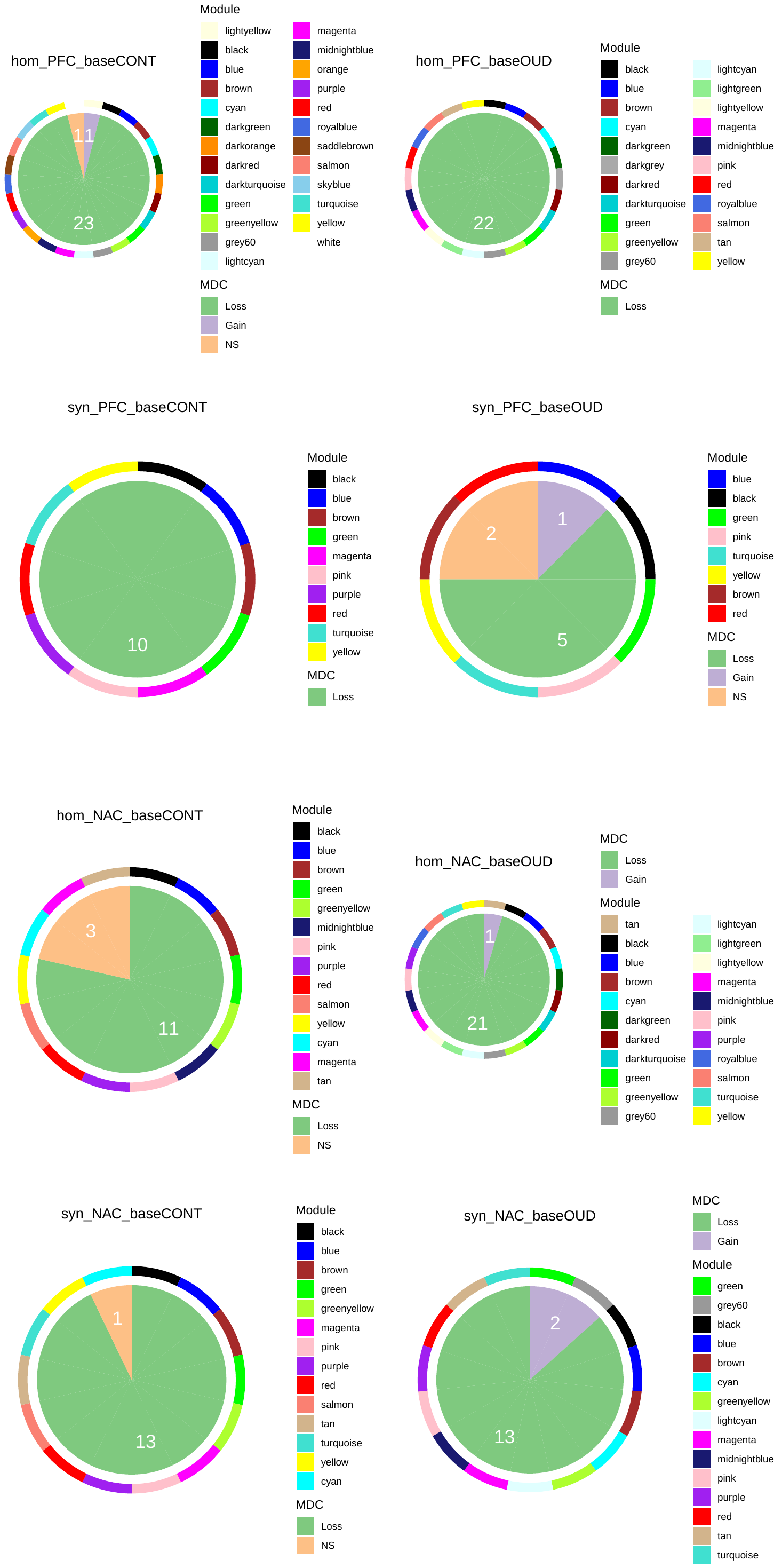

**Figure S11. Modulate differential connectivity of WGCNA modules.**

**Table S1. Subject information.** Postmortem brains were collected from people diagnosed with OUD and from matched unaffected comparison subjects. Subjects were matched on age, sex, postmortem interval (PMI), brain pH, and RNA integrity (RIN). All brains were provided by the brain bank in the Department of Psychiatry at the University of Pittsburgh School of Medicine.

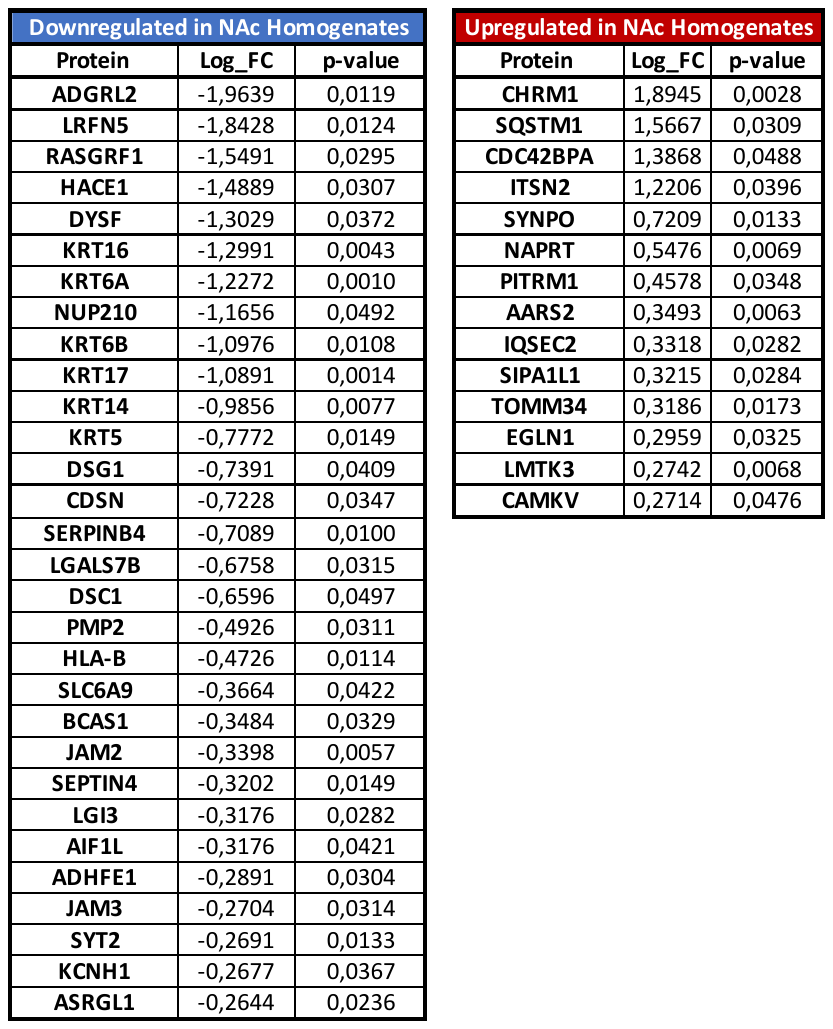

**Table S2. Significant DE proteins in NAc homogenates.** Differentially expressed proteins in tissue homogenates from NAc of subjects with OUD compared to unaffected subjects. We identified 43 DE (p<0.05 and logFC±0.26) proteins (14 upregulated and 29 downregulated).

**Table S3. Full protein expression list in NAc homogenates.** Full list of proteins detected in tissue homogenates from NAc, and levels of expression in OUD compared to unaffected subjects.

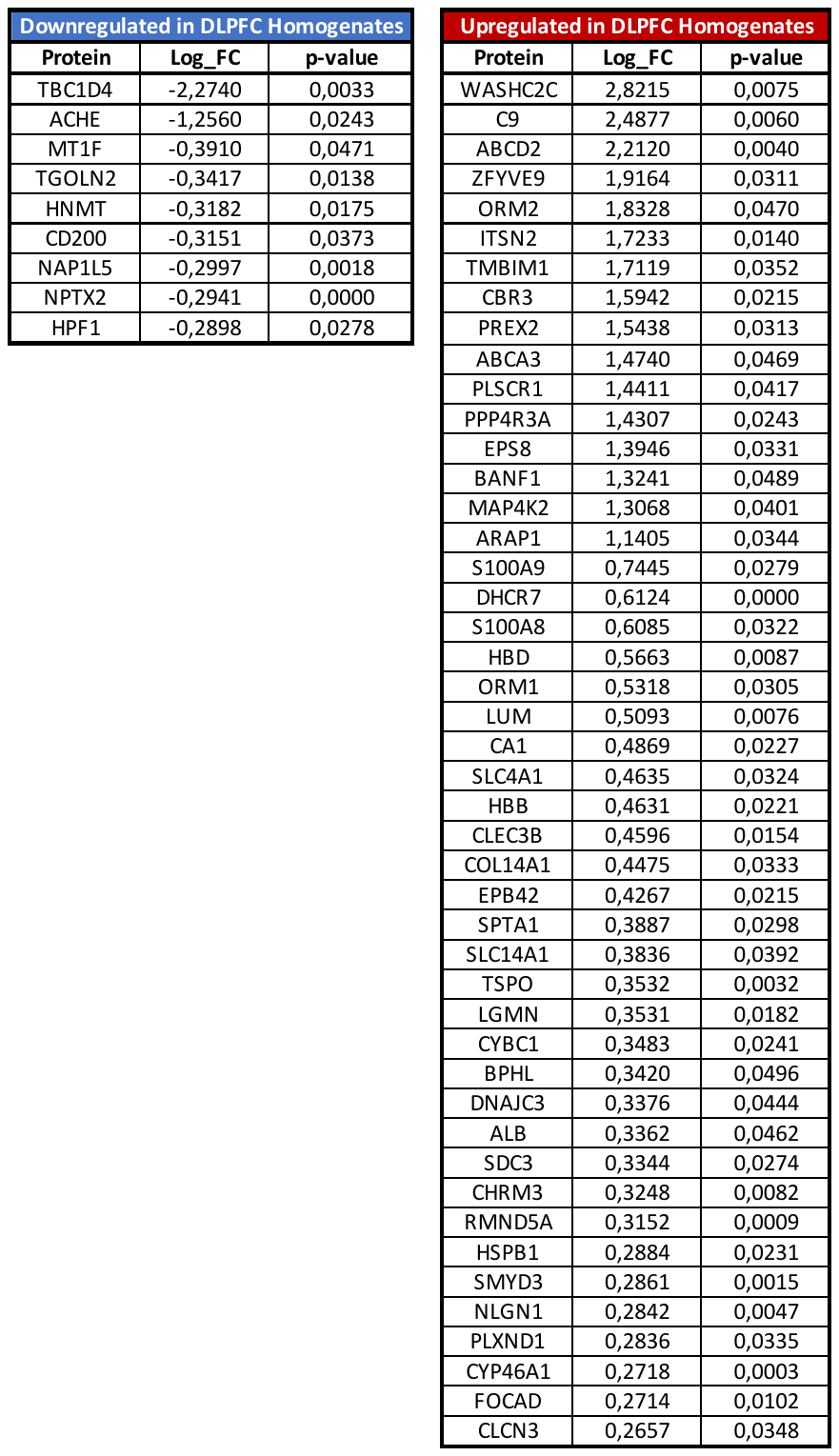

**Table S4. Significant DE proteins in DLPFC homogenates.** Differentially expressed proteins in tissue homogenates from DLPFC of subjects with OUD compared to unaffected subjects. We identified 55 DE proteins (p<0.05 and logFC±0.26), with a majority being upregulated in OUD (46 upregulated and 9 downregulated).

**Table S5. Full protein expression list in DLPFC homogenates.** Full list of proteins detected in tissue homogenates from DLPFC, and levels of expression in OUD compared to unaffected subjects.

**Table S6. Pathway enrichment in NAc and DLPFC homogenates and synaptosomes.**

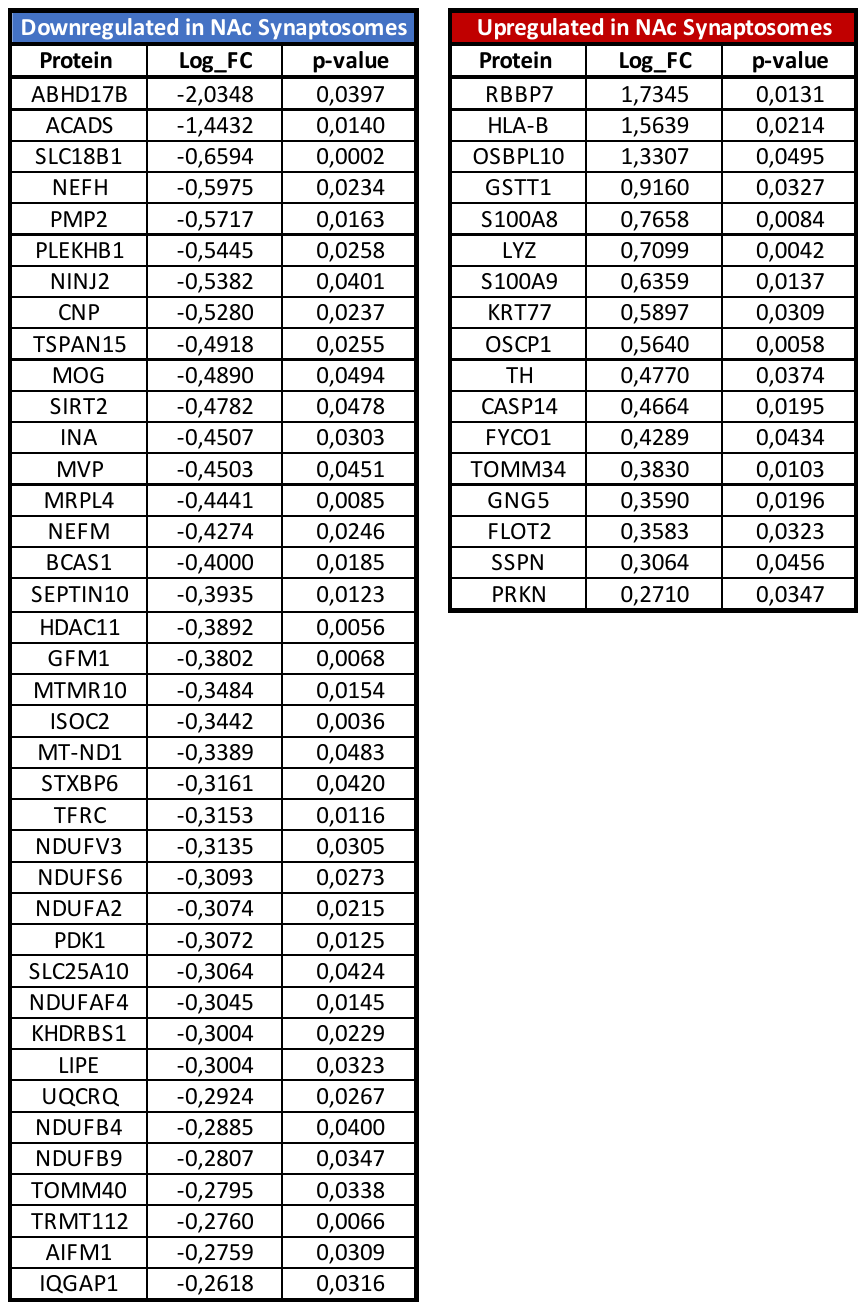

**Table S7. Significant DE proteins in NAc synaptosomes.** Differentially expressed proteins in NAc Synaptosomes of subjects with OUD compared to unaffected subjects. We identified 56 DE proteins (p<0.05 and logFC±0.26), with a majority being upregulated in OUD (17 upregulated and 39 downregulated). DE, Differentially Expressed; NAc Nucleus Accumbens.

**Table S8. Full protein expression list in NAc synaptosomes.** Full list of proteins detected in synaptosomes from NAc and levels of expression in OUD compared to unaffected subjects.

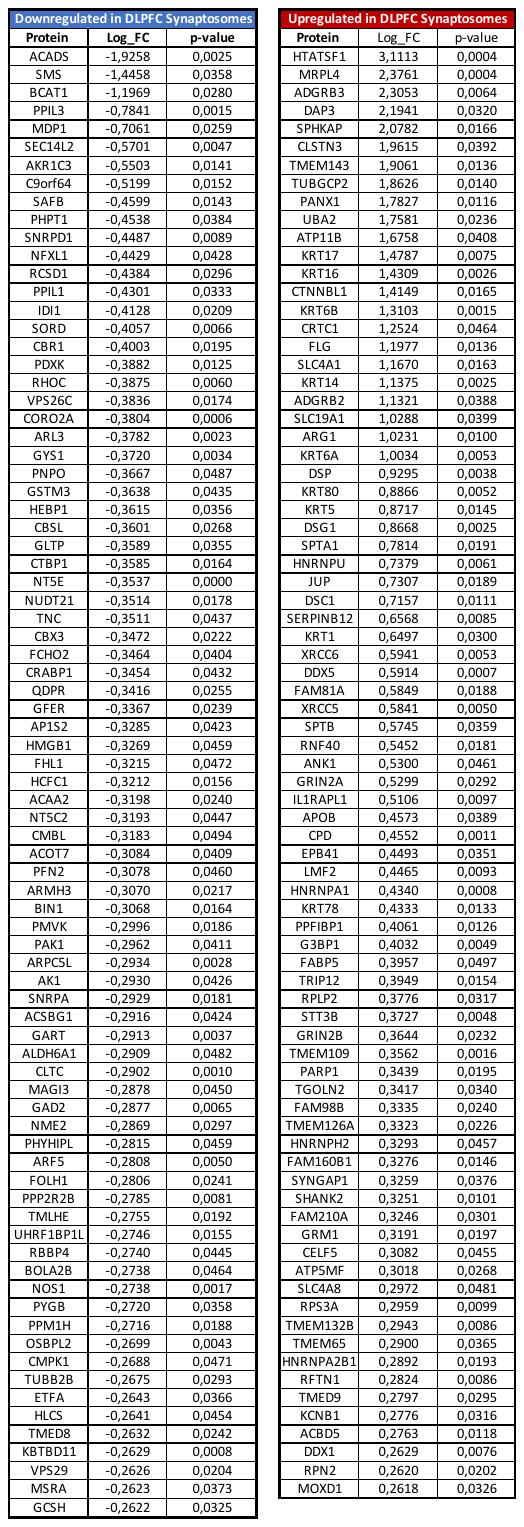

**Table S9. Significant DE proteins in DLPFC synaptosomes.** Differentially expressed proteins in DLPFC synaptosomes of subjects with OUD compared to unaffected subjects. We identified 161 DE proteins (p<0.05 and logFC±0.26), with a majority being upregulated in OUD (80 upregulated and 81 downregulated). DE, Differentially Expressed; NAc Nucleus Accumbens.

**Table S10. Full protein expression list in DLPFC synaptosomes.** Full list of proteins detected in tissue synaptosomes from DLPFC and levels of expression in OUD compared to unaffected subjects.

**Table S11. Protein enrichment in NAc synaptosomes.**

**Table S12. Protein enrichment in DLPFC synaptosomes.**

**Table S13. Rhythmic proteins in NAc homogenates in unaffected subjects.** Analysis of rhythms (amplitude, phase, peak) in NAc homogenates from unaffected subjects, generated by plotting protein expression across time of day.

**Table S14. Rhythmic proteins in NAc homogenates in OUD subjects.** Analysis of rhythms (amplitude, phase, peak) in NAc homogenates from OUD subjects, generated by plotting protein expression across time of day.

**Table S15. Rhythmic proteins in DLPFC homogenates in unaffected subjects.** Analysis of rhythms (amplitude, phase, peak) in DLPFC homogenates from unaffected subjects, generated by plotting protein expression across time of day.

**Table S16. Rhythmic proteins in DLPFC homogenates in OUD subjects.** Analysis of rhythms (amplitude, phase, peak) in DLPFC homogenates from OUD subjects, generated by plotting protein expression across time of day.

**Table S17. Rhythmic proteins in NAc synaptosomes in unaffected subjects.** Analysis of rhythms (amplitude, phase, peak) in NAc synaptosomes from Unaffected subjects, generated by plotting protein expression across time of day.

**Table S18. Rhythmic proteins in NAc synaptosomes in OUD subjects.** Analysis of rhythms (amplitude, phase, peak) in NAc Synaptosomes from OUD subjects, generated by plotting protein expression across time of day.

**Table S19. Rhythmic proteins in DLPFC synaptosomes in unaffected subjects.** Analysis of rhythms (amplitude, phase, peak) in DLPFC synaptosomes from Unaffected subjects, generated by plotting protein expression across time of day.

**Table S20. Rhythmic proteins in DLPFC synaptosomes in OUD subjects.** Analysis of rhythms (amplitude, phase, peak) in DLPFC synaptosomes from OUD subjects, generated by plotting protein expression across time of day.

**Table S21. Gain and loss of protein rhythms in NAc homogenates.** Analysis of gain and loss of rhythms via comparison of amplitude, phase and peak of rhythmic proteins in NAc homogenates.

**Table S22. Gain and loss of protein rhythms in DLPFC homogenates.** Analysis of gain and loss of rhythms via comparison of amplitude, phase and peak of rhythmic proteins in DLPFC homogenates.

**Table S23. Gain and loss of protein rhythms in NAc synaptosomes.** Analysis of gain and loss of rhythms via comparison of amplitude, phase and peak of rhythmic proteins in NAc synaptosomes.

**Table S24. Gain and loss of protein rhythms in DLPFC synaptosomes.** Analysis of gain and loss of rhythms via comparison of amplitude, phase and peak of rhythmic proteins in DLPFC synaptosomes.

**Table S25. WGCNA module assignments.** Weight Gene Correlation Network Analysis (WGCNA) module assignment in NAc and DLPFC homogenates and synaptosomes.

**Table S26. MDC analysis.** Module Differential Connectivity (MDC) analysis in NAc and DLPFC homogenates and synaptosomes.
